# Dynamic Brain-Body Coupling of Breath-by-Breath O_2_-CO_2_ Exchange Ratio with Resting State Cerebral Hemodynamic Fluctuations

**DOI:** 10.1101/843482

**Authors:** Suk-tak Chan, Karleyton C. Evans, Tian-yue Song, Juliette Selb, Andre van der Kouwe, Bruce R. Rosen, Yong-ping Zheng, Andrew C. Ahn, Kenneth Kwong

**Affiliations:** Athinoula A. Martinos Center for Biomedical Imaging, Department of Radiology, Massachusetts General Hospital, MA; Biogen Inc., Cambridge, MA; Department of Biomedical Engineering, The Hong Kong Polytechnic University, Hong Kong

**Keywords:** Resting State, Blood oxygenation level dependent, Functional magnetic resonance imaging, Breath-by-breath O_2_-CO_2_ exchange ratio, Cerebral blood flow velocity, Transcranial Doppler sonography

## Abstract

The origin of low frequency cerebral hemodynamic fluctuations (CHF) in resting state remains unknown. Here we studied the contribution of respiratory gas exchange (RGE) metrics to CHF during spontaneous breathing. RGE metrics include the breath-by-breath changes of partial pressure of oxygen (ΔPO_2_) and carbon dioxide (ΔPCO_2_) between end inspiration and end expiration, and their ratio breath-by-breath O_2_-CO_2_ exchange ratio (bER). We used transcranial Doppler sonography to evaluate CHF changes during spontaneous breathing by measuring the cerebral blood flow velocity (CBFv) in the middle cerebral arteries. The regional CHF changes during spontaneous breathing were mapped with blood oxygenation level dependent (BOLD) signal changes using functional magnetic resonance imaging (fMRI) technique. We found that prominent oscillations with periods of 0.5 to 2 minutes characterized ΔPO_2_, ΔPCO_2_ and bER. The oscillations of bER were coherent with CHF during spontaneous breathing at the frequency range of 0.008-0.03Hz which is consistent with the low frequency resting state CHF. CHF had strong correlation with bER, followed by ΔPO_2_ and then by ΔPCO_2_. Brain regions with the strongest bER-CHF coupling overlapped with many areas of default mode network. Although the physiological mechanisms underlying the strong correlation between bER and CHF are not completely understood, our findings suggest the contribution of bER to low frequency resting state CHF. It also provides a novel insight of brain-body interaction via CHF and oscillations of RGE metrics.

## INTRODUCTION

Low frequency components (below 0.05 Hz) in the cerebral hemodynamic fluctuations (CHF) are critical for characterizing functional connectivity of cerebral resting state networks including the default mode network (DMN) (Biswal et al., 1995; Greicius et al., 2003). The feature of CHF most useful for the evaluation of cerebral connectivity is the pattern of low frequency fluctuation and not the time-averaged hemodynamic value. The origin of these low frequencies of CHF is still not clear and many physiological candidates of cardiac and respiratory origin had been proposed to contribute in this frequency bandwidth (Birn et al., 2006; Chang and Glover, 2009; Chang et al., 2013; Wise et al., 2004). They include respiratory variation related fluctuations (∼0.03Hz) (Birn et al., 2006; Chang and Glover, 2009), heart rate variability (0.05-0.15Hz) (Chang et al., 2013) and end-tidal carbon dioxide fluctuations (0-0.05Hz) (Wise et al., 2004). These findings in general suggest an interaction between resting state CHF and elements of ventilation.

The objective in our study was to investigate whether respiratory gas exchange (RGE) metrics during spontaneous breathing at rest can also be candidates contributing to low frequencies of CHF. RGE involves a process of removing carbon dioxide (CO_2_) from blood and replenishing oxygen (O_2_). The RGE metrics we investigated were breath-by-breath changes in partial pressure of CO_2_ and O_2_, and O_2_-CO_2_ exchange ratio. Oscillations in end-tidal partial pressures of O_2_ and CO_2_ (P_ET_O_2_ and P_ET_CO_2_) had been reported to be in the low frequency range below 0.05 Hz in humans (Lenfant, 1967; Van den Aardweg and Karemaker, 2002). The source of such oscillations may be attributed to the feedback control of ventilation via chemoreceptor activities (Van den Aardweg and Karemaker, 2002). Early study by Lahiri et al. (1978) reported that even under normoxic conditions (partial pressure of O_2_ at 100 mmHg or above) both arterial partial pressure of O_2_ (PO_2_) and partial pressure of CO_2_ (PCO_2_) in cats interacted synergistically in the feedback control loop of ventilation with chemoreceptor reflexes. Studies on animal models showed that the feedback loops involved interaction between central respiratory chemoreceptors in the brainstem (Guyenet et al., 2010; Nattie and Li, 2012) and peripheral respiratory chemoreceptors at the carotid body (Daristotle et al., 1987; Lahiri and DeLaney, 1975b).

According to concept of homeostasis, homeostatic regulatory system is formed with arterial PO_2_ and PCO_2_ as regulated variables, peripheral and central chemoreceptors as sensors, brain stem as control center, and diaphragm and respiratory muscles as effectors, to optimize systemic blood gases. While the detailed mechanisms between the change in central chemoreceptor activities and the change in CBF are topics of on-going research (Cummins et al., 2019; Kumar and Prabhakar, 2012; Rocha and Branco, 1998), arterial PO_2_ and PCO_2_ which are sensed by chemoreceptors are maintained within a range (or fluctuate) around the physiological ‘set point’ (i.e. mean) by the feedback control of diaphragm and respiratory muscles for ventilation in the homeostatic regulatory system. Spontaneous breathing is therefore part of a vital homeostatic process to optimize the systemic blood gases which can presumably regulate in turn cerebral blood flow (CBF) and O_2_ delivery to the brain that is compatible with the viability of individual (Guyton, 1966; Modell et al., 2015). We hypothesize that such homeostatic fluctuations of arterial PO_2_ or PCO_2_ interact synergistically and may contribute to fluctuations of CHF during spontaneous breathing at rest.

Given the dynamic changes in PO_2_ and PCO_2_ breath-by-breath in humans, we studied RGE metrics as surrogates of arterial blood gases with the recognition of the difference between respiratory metrics and blood gases. We studied the interaction between RGE metrics and CHF measured with transcranial Doppler sonography (TCD) and functional magnetic resonance imaging (fMRI). TCD has an advantage of acquiring data at high temporal resolution (∼100 Hz) without much concern on high frequency cerebrovascular signal being aliased into the low frequency range. The cerebral blood flow velocities (CBFv) were measured by TCD in the middle cerebral arteries (MCA) which supply most parts of the brain. The application of TCD allows the subjects to have measurements in upright seated position. Although TCD offers high temporal resolution to evaluate CHF, it does not provide regional information. Regional mapping of spontaneous CHF was carried out instead with blood oxygen level-dependent (BOLD) signal changes measured by fMRI. We used BOLD-fMRI instead of arterial spin labeling (ASL) in MRI perfusion for two major reasons. ASL perfusion-related signal has low contrast to noise ratio in comparison with BOLD signal especially in white matter area. ASL image acquisition at a temporal resolution of 4 seconds also under-samples the spontaneous fluctuations within respiratory cycle of 4-6 seconds, which hampers the dynamic analyses between CHF and RGE metrics. While CBFv and BOLD signals are not equivalent surrogates for cerebral blood flow, combining their strengths would be powerful to study the temporal and spatial features in the association between RGE and CHF.

In addition to the more commonly used P_ET_O_2_ and P_ET_CO_2_, we selected to investigate in this manuscript the RGE metrics which included changes in partial pressure of O_2_ (ΔPO_2_) and of CO_2_ (ΔPCO_2_) between end inspiration and end expiration and their ratio (ΔPO_2_/ΔPCO_2_) which we named breath-by-breath exchange ratio (bER). We preferred to use the terms ΔPO_2_ (defined as expired PCO_2_ – inspired PCO_2_) and ΔPCO_2_ (defined as inspired PO_2_ – expired PO_2_) over P_ET_O_2_ and P_ET_CO_2_ partly to take into account any variation in the inspired ambient air. A more useful reason was that ΔPCO_2_ and ΔPO_2_ share the same polarity of changes and similar dynamic partial pressure range of about 30 to 50mm Hg while P_ET_CO_2_ and P_ET_O_2_ oscillate out of phase and have a very different dynamic range. ΔPCO_2_ and P_ET_CO_2_ are basically interchangeable due to the low level of ambient CO_2_. Unlike P_ET_O_2_, ΔPO_2_ is related to the actual partial pressure of O_2_ being utilized by the whole body. More importantly, ΔPO_2_ and not P_ET_O_2_ is the designated term used to study RGE in the alveolar air equation (Fenn et al., 1946; Ferretti, 2015). While ΔPCO_2_ and P_ET_CO_2_ are interchangeable, P_ET_O_2_ and ΔPO_2_ differ by a constant if ambient O_2_ remains steady.

bER is of special interest because as a ratio it factors out ventilatory volume fluctuations common to ΔPO_2_ and ΔPCO_2_, reducing the unwanted effects of ventilatory volume due to factors like isolated deep breaths on the study of the interaction between bER and CHF. Being a ratio of two gas metrics, the bER formulation took into account the natural interdependence of ΔPO_2_ and ΔPCO_2_ in the RGE process and in chemoreceptor activities mentioned above. bER is a breath-by-breath dynamic form related to the steady-state respiratory exchange ratio (RER) described in the alveolar gas equation introduced by Fenn et al. (1946); time-averaged value of bER over minutes is mathematically equivalent to the reciprocal of RER. RER has been used to evaluate resting systemic metabolic rate (Bain et al., 2012; Nishi, 1981; Weir, 1949). There are also technical differences between RER used in the literature and bER we used in this study. Traditionally, RER is derived by measuring the respiratory flow and the expired gases collected in Douglas bag connected to a closed circuit over several minutes. In our study, bER was derived by measuring the inspired and expired gases with a nasal tubing at each breath. We hypothesized that the fluctuations of bER were not just random noises but were related to the fluctuations of systemic ΔPO_2_ and ΔPCO_2_ which came under the influences of systemic metabolism and/or of ambient gases as well as the feedback from the chemoreceptors.

In the present study, we evaluated the brain-body interaction between resting state CHF and the dynamic fluctuations of ΔPO_2_, ΔPCO_2_ and bER. Before we correlated the changes of RGE metrics with CBFv and BOLD signal changes during spontaneous breathing, we examined the correlations among these respiratory parameters measured in both TCD and MRI sessions. In the TCD study, we measured simple correlation of CBFv in the MCA with RGE metrics of bER, ΔPO_2_ and ΔPCO_2_. We also examined the temporal features and frequency characteristics of these RGE metrics and their coherence with CBFv. In the fMRI study, we mapped the association between regional BOLD signal changes and RGE metrics of bER, ΔPO_2_ and ΔPCO_2_. Brain regions showing significant association between regional BOLD signal changes and RGE metrics were then compared with regions within the default mode network (DMN) outlined by the resting state connectivity analysis with the seed at the left precuneus. DMN encompasses brain regions responsible for the constant background activities at rest, showing higher metabolic and hemodynamic level than the other parts of the cortex (Raichle et al., 2001). The temporal features and frequency characteristics of these RGE metrics and their coherence with BOLD signal changes in the regions within DMN were also examined. To verify if the potential coherence between CHF and RGE metrics were resulting from respiratory variability, the respiratory metric of respiration volume per unit time (RVT) was also derived for comparison. The investigation of the association between RGE metrics and CHF, in addition to offering a physiological model to characterize the contribution of gas exchange elements in the low frequency resting state fluctuations, would also provide new directions to the study of brain-body interaction. The potential of using the interaction between CHF and RGE metrics to evaluate cerebrovascular reactivity (CVR) to elements in spontaneous breathing was also discussed.

## MATERIALS AND METHOD

### Participants

Twenty-two volunteers aged from 19 to 48 years (mean age = 30.5 years, SD = 9.1 years, 14 males and 8 females) were included. Eleven of them participated in both TCD and MRI sessions, while the remaining participated in either one of the sessions. Subject demographics were shown in Table 1. All of them were recruited by e-mail and poster placement within the Partners hospital network. They were screened to exclude neurological, mental and medical disorders and drug abuse. TCD and MRI scanning were performed in the Athinoula A. Martinos Center for Biomedical Imaging at the Massachusetts General Hospital of Partners HealthCare. All the experimental procedures were explained to the subjects, and signed informed consent was obtained prior to participation in the study. All components of this study were performed in compliance with the Declaration of Helsinki and all procedures were approved by Partners Human Research Committee.

**Table 1.** Subject demographics and their participation in the TCD and MRI sessions.

| **Subjects** | **Gender** | **Age** | **TCD** | **MRI** |
| --- | --- | --- | --- | --- |
| s1 | M | 35 | - | √ |
| s2 | M | 48 | - | √ |
| s3 | M | 22 | - | √ |
| s4 | M | 46 | - | √ |
| s5 | M | 32 | - | √ |
| s6 | M | 29 | - | √ |
| s7 | F | 45 | - | √ |
| s8 | M | 19 | - | √ |
| s9 | M | 20 | - | √ |
| s10 | M | 38 | √ | √ |
| s11 | M | 28 | √ | √ |
| s12 | M | 27 | √ | √ |
| s13 | M | 32 | √ | √ |
| s14 | M | 22 | √ | √ |
| s15 | M | 32 | √ | √ |
| s16 | F | 26 | √ | √ |
| s17 | F | 27 | √ | √ |
| s18 | F | 47 | √ | √ |
| s19 | F | 25 | √ | √ |
| s20 | F | 23 | √ | √ |
| s21 | F | 25 | √ | - |
| s22 | F | 23 | √ | - |

Our study was divided into two parts: Part I and Part II. In Part I, we aimed to correlate the RGE metrics including bER, ΔPCO_2_ and ΔPO_2_ with CBFv in MCAs as CHF in the major cerebral arterial supply during spontaneous breathing measured by TCD. We also examined the temporal features and frequency characteristics of these RGE metrics and their coherence with CBFv. Thirteen subjects participated in TCD sessions. In Part II, we aimed to map the association between these RGE metrics (bER, ΔPCO_2_ and ΔPO_2_) and BOLD signal changes as regional CHF during spontaneous breathing. Twenty subjects participated in MRI sessions. Before we correlated the changes of RGE with CBFv and BOLD signal changes, we examined the correlations among the respiratory metrics (bER, ΔPCO_2_ and ΔPO_2_) acquired in both TCD and MRI sessions.

### Part 1: TCD

#### Transcranial Doppler scanning

Before the study of blood flow velocity in intracranial arteries, subject was allowed to rest at least 20-30 minutes for hemodynamic stabilization. The blood pressure measured in the subject was within the normal range (Chobanian et al., 2003). With the subject in an upright seated position, a dual probe setting with 2MHz transducers in conjunction with TCD system (Delicate EMS-9U, Shenzhen, China) was used for simultaneous recording of CBFv in the MCA on both left and right sides while the subject was at rest. Two transducers were attached onto the left and right temporal bone windows by velcro. The depth of the Doppler samples was confined to the M1 segment, which is at the main stem of the MCA, for all the subjects.

A white crosshair in black background was presented visually to the subject by a computer using the software Eprime Professional 2.0 (Psychology Software Tools, Inc., Pittsburgh, USA) to fixate their eye movement. The total duration of the CBFv data acquisition lasted 10 minutes.

Physiological changes including PCO_2_, PO_2_, electrocardiogram (ECG) and peripheral blood pressure were measured simultaneously with TCD acquisition. A small nasal tubing was placed at the subject’s nostril to sample PCO_2_ and PO_2_ via gas analyzers (Capstar-100, Oxystar-100, CWE, Inc., PA, USA) after calibrating to the barometric pressure of the day of TCD session and correcting for vapor pressure. Small nasal tubing was chosen to sample PCO_2_ and PO_2_ because of several reasons. In the current study, we standardized all the physiological set-up in both TCD and MRI sessions. We did not measure PCO_2_ and PO_2_ from a facemask connected to a closed breathing circuit with non-rebreathing valves or from the expired gases collected in Douglas bag because of the space constraint of the MRI head coil and the slow sampling of gases in the units of minutes respectively. Sampling gases with small nasal tubing has an advantage that the subjects breathed normally through the nose. In the gas sampling circuit, the same gas sample volume was used in both CO_2_ and O_2_ analyzers at the same gas sampling flow rate. The CBFv and physiological measurements were synchronized using trigger signals from E-prime. All the CBFv time series and physiological recordings were stored for offline data analysis.

### Part 2: MRI

#### MRI acquisition

MRI brain scanning was performed on a 3-Tesla scanner (Siemens Medical, Erlangen, Germany). The head was immobilized in a standard head coil with foam pads. The following whole brain MRI datasets were acquired on each subject: 1) standard high-resolution sagittal images acquired with volumetric T1-weighted 3D-MEMPRAGE (TR=2530ms, TE=1.74ms/3.6ms/5.46ms/7.32ms, flip angle=7°, FOV=256×256mm, matrix=256×256, slice thickness=1mm); 2) BOLD-fMRI images acquired with gradient-echo echo planar imaging (EPI) sequence (TR=1450ms, TE=30ms, flip angle=90°, FOV=220×220mm, matrix=64×64, thickness=5mm, slice gap=1mm) while the subject was at rest. The visual presentation of crosshair, the physiological set-up for sampling of PCO_2_ and PO_2_ in the MRI session were the same as those used in the TCD session. The gas analyzers were again calibrated to the barometric pressure of the day of MRI session and corrected for vapor pressure. ECG was measured using Siemens physiological monitoring unit (Siemens Medical, Erlangen, Germany). Physiological changes including PCO_2_, PO_2_, ECG and respiration were measured simultaneously with MRI acquisition. All the physiological measurements including those of PCO_2_, PO_2_ and ECG were synchronized using trigger signals from the MRI scanner. BOLD-fMRI images and physiological recordings were stored for offline data analysis.

#### Exogenous CO_2_ challenge

Ten out of 20 subjects had additional exogenous CO_2_ challenge in the MRI sessions. Given that there is significant inter-individual variance in resting P_ET_CO_2_ (West, 1992), resting P_ET_CO_2_ was assessed in subjects via calibrated capnograph before the exogenous CO_2_ challenge. Subject wore nose-clip and breathed through a mouth-piece on an MRI-compatible circuit designed to maintain the P_ET_CO_2_ within ± 1-2 mmHg of target P_ET_CO_2_ (Banzett et al., 2000; McKay et al., 2003). The fraction of inspired carbon dioxide was adjusted to produce steady-state conditions of normocapnia and mild hypercapnia (4-8 mmHg above the subject’s resting P_ET_CO_2_). The CO_2_ challenge paradigm consisted of 2 consecutive phases (normocapnia and mild hypercapnia) repeating 6 times with 3 epochs of 4 mmHg increase and 3 epochs of 8 mmHg increase of P_ET_CO_2_. The normocapnia phase lasted 60-90 seconds, while the mild hypercapnia phase lasted 30 seconds. The total duration of the exogenous CO_2_ hypercapnic challenge lasted 10 minutes.

When the subject had exogenous CO_2_ challenge in MRI session, BOLD-fMRI images were acquired using the same EPI sequence for resting state. The PCO_2_ and PO_2_ were sampled through the air filter connected with the mouthpiece and the sampled gases were measured by calibrated gas analyzers. The respiratory flow was measured with respiratory flow head (MTL300L, ADInstruments, Inc., CO, USA) on the breathing circuit via calibrated spirometer (FE141, ADInstruments, Inc., CO, USA). The physiological measurements were synchronized with MRI images using trigger signals from MRI scanner. All the BOLD-fMRI images and physiological recordings were stored for offline data analysis.

## DATA ANALYSIS

### Processing of physiological data

The physiological data from both TCD and MRI sessions were analyzed using Matlab R2014a (Mathworks, Inc., Natick, MA, USA). Technical delay of PCO_2_ and PO_2_ was corrected by cross-correlating the time courses of PCO_2_ and PO_2_ with respiratory phases determined from the artifactual displacement due to chest excursion on ECG time courses in the TCD sessions, with the respiratory phases from respiratory bellow or respiratory flow measured with respiratory flow head via spirometer in the MRI sessions.

End inspiration (I) and end expiration (E) were defined on the time courses of PO_2_ and PCO_2_. The breath-by-breath P_ET_CO_2_ and P_ET_O_2_ were extracted at the end expiration of PCO_2_ and PO_2_ time courses respectively. ΔPO_2_ is defined as (inspired PO_2_ – expired PO_2_) and ΔPCO_2_ is defined as (expired PCO_2_ – inspired PCO_2_). Breath-by-breath O_2_-CO_2_ exchange ratio (bER) is defined as the ratio of ΔPO_2_ to ΔPCO_2_ measured between end inspiration and end expiration at each breath, whereas RER at steady state from alveolar air equation introduced by Fenn et al. (1946) is formulated as the ratio of ΔPCO_2_ to ΔPO_2_ over minutes. The product of ΔPO_2_ and ΔPCO_2_ was not used to evaluate the interaction with CHF because the effects from fluctuations due to ventilation would be exacerbated in ΔPO_2_×ΔPCO_2_ (Figure S1 in Supplementary materials).

Simple correlation analyses were applied on the time series of RGE metrics (bER, ΔPCO_2_, and ΔPO_2_) in pairs. Correlation was considered significant at p<0.05.

### Part 1

#### Preprocessing of CBFv data

The CBFv data were analyzed using Matlab R2014a (Mathworks, Inc., Natick, MA, USA). A median filter was applied to the data to reduce artifactual spikes. Beat-by-beat systolic peaks and end-diastolic troughs were determined using custom Matlab function and corrected on the graphical user interface incorporated in the function. Systolic peaks and diastolic troughs of cardiac cycles on the CBFv time courses showing persistent artifacts were excluded in the following analysis. TCD data in both left and right MCAs were acquired on 13 subjects. One of the 13 TCD datasets had persistent artifacts in over one-third of the CBFv time courses acquired in the LMCA and another one had persistent artifacts in CBFv data acquired in RMCA. The CBFv time courses in the LMCA of those particular TCD runs were excluded in further analysis, while the CBFv time courses without persistent artifacts in the contralateral MCA were retained. Time courses of mean CBFv was derived by integrating the CBFv over each cardiac cycle. In order to reduce the large inter-individual variations of absolute blood flow velocities (Aaslid et al., 1982; Ringelstein et al., 1990) and to remove the dependence of insonation angle (Deppe et al., 1997), the percent change of CBFv (ΔCBFv) of the left and right MCAs relative to baseline value was derived. The mean CBFv for a period of 30 seconds at the beginning of the time courses was chosen as the baseline.

#### Correlation analyses between **Δ**CBFv and RGE metrics

The time courses of ΔCBFv were correlated with those of bER, ΔPCO_2_ and ΔPO_2_ separately. The correlation indicated by Pearson’s correlation coefficient was considered significant at p<0.05. Fisher Z-transformation was used to transform Pearson’s correlation coefficients to Fisher’s z scores for group analysis. Paired t-tests were used to compare the Fisher’s z scores representing the correlation between ΔCBFv and bER with those indicating the correlation between ΔCBFv and other physiological parameters besides bER. Differences were considered to be significant at p<0.05.

#### Dynamic analysis of coherence between **Δ**CBFv and RGE metrics as function of time and frequency

Wavelet transform coherence (WTC) was employed to demonstrate the dynamic interaction between ΔCBFv and RGE metrics (Figure S2 in Supplementary materials) in the time-frequency domain. WTC is a method for analyzing the coherence and phase lag between two time courses as a function of both time and frequency (Grinsted et al., 2004; Torrence and Compo, 1998). The temporal and phase information of WTC have been used to map dynamic connectivity of the brain regions related to heart rate changes (Keilholz et al., 2018). It is therefore well suited to investigating dynamic changes in coupling between the time courses of ΔCBFv and RGE metrics including bER, ΔPCO_2_, and ΔPO_2_, as well as the phase lag of ΔCBFv to bER, ΔPCO_2_ and ΔPO_2_. We used the Matlab wavelet cross-spectrum toolbox developed by Grinsted et al. (2004). Squared wavelet coherence between the time courses of each RGE metric and ΔCBFv is separately plotted with x-axis as time and y-axis as scale which has been converted to its equivalent Fourier period. An example of squared wavelet coherence between bER and ΔCBFv in right MCA from a representative subject during spontaneous breathing is shown in Figure S2B (Supplementary materials). The magnitude of wavelet transform coherence ranged between 0 and 1 that can be conceptualized as a localized correlation coefficient in time and frequency space (Grinsted et al., 2004). The phase angle between the two time courses, with bER leading ΔCBFv, at particular samples of the time-frequency plane is indicated by an arrow: a rightward pointing arrow indicates that the time courses are in phase, or positively correlation (ϕ=0); a leftward pointing arrow indicates anti-correlation (ϕ=π), and the downward and upward pointing arrows indicate phase angles of π/2 and -π/2 relative to ϕ=0, respectively. Areas inside the ‘cone of influence’, which are locations in the time-frequency plane where edge effects give rise to lower confidence in the computed values, are shown in faded color outside of the conical contour. The statistical significance level of the wavelet coherence is estimated using Monte Carlo method and the 5% significance level against red noise is shown as thick contour in the squared wavelet coherence plot. The wavelet coherence magnitudes and phases bounded by thick contour outside the cone of influence are considered significant.

Time-averaged coherence is defined as the total significant coherence at each scale of Fourier periods (converted into frequency) where the wavelet coherence magnitude exceeded 95% significance level, normalized by the maximum possible coherence outside the cone of influence at that particular scale (Figure S2C in Supplementary materials). It is interpreted in the similar way as the coherence in the transfer function analysis which has been used in cerebral autoregulation study (Zhang et al., 1998).

Interpreting the time-averaged coherence irrespective of phase lag raised the question that the coherence with positive correlation between two time courses (at phase lag of 0±π/2) and the coherence with negative correlation (at phase lag of π±π/2) might be mixed together. Therefore, mean time-averaged coherence at the phase lags of 0±π/2 and π±π/2 were separately averaged across all the subjects who participated in the TCD sessions to explore the Fourier periods/frequency bandwidths that oscillations of ΔCBFv were in synchrony with the time courses of each RGE metric (bER, ΔPCO_2_ and ΔPO_2_) when they were at rest.

### Part 2

#### Preprocessing of BOLD-fMRI data

All the BOLD-fMRI data were imported into the software Analysis of Functional NeuroImage (AFNI) (Cox, 1996) (National Institute of Mental Health, http://afni.nimh.nih.gov) for time-shift correction, motion correction, normalization and detrending. The first 12 volumes in the first 12 time points of each functional dataset, collected before equilibrium magnetization was reached, were discarded. Each functional dataset was corrected for slice timing, motion-corrected and co-registered to the first image of the first functional dataset using three-dimensional volume registration. It was then normalized to its mean intensity value across the time-series. Voxels located within the ventricles and outside the brain defined in the parcellated brain volume using FreeSurfer (Dale et al., 1999; Fischl et al., 1999) (MGH/MIT/HMS Athinoula A. Martinos Center for Biomedial Imaging, Boston, http://surfer.nmr.mgh.harvard.edu) were excluded from the following analyses of functional images. The time-series of each voxel in the normalized functional dataset was detrended with the 5^th^ order of polynomials to remove the low drift frequency. Individual brain volumes with time series of percent BOLD signal changes (ΔBOLD) were derived.

#### Linear regression for the association between **Δ**BOLD and RGE metrics in individual subject

Linear regression analysis was used to evaluate the association between ΔBOLD, bER, ΔPO_2_ and ΔPCO_2_ for each subject. For each voxel of the preprocessed brain volume, time course of ΔBOLD was regressed on bER, ΔPO_2_ and ΔPCO_2_, which served as a regressor in separate linear regression analyses. Regression coefficient beta (β) value was defined as the percent BOLD signal changes per unit change of the regressor. Individual subject brain volumes with β magnitudes were registered onto their own anatomical scans and transformed to the standardized space of Talairach and Tournoux (1988). Monte Carlo simulation was used to correct for multiple comparisons (Gold et al., 1998). In order to protect against type I error, individual voxel probability threshold of p<0.005 was held to correct the overall significance level to α<0.05. Based upon a Monte Carlo simulation with 2000 iteration processed with ClustSim program (Ward, 1997), it was estimated that a 476mm^3^ contiguous volume would provide the significance level α<0.05, which met the overall corrected threshold of p<0.05.

#### Group region-of-interest (ROI) analysis for the association between **Δ**BOLD and RGE metrics

For each subject who participated in MRI scanning, β magnitude values derived by regressing ΔBOLD on bER (β_bER_), ΔBOLD on ΔPO_2_ (β_ΔPO2_) and ΔBOLD on ΔPCO_2_ (β_ΔPCO2_) were separately averaged in each of the 160 brain regions parcellated by the software FreeSurfer. One-sample t-tests were used to analyze regional β_bER_, β_ΔPO2_ and β_ΔPCO2_ in subject group. False discovery rate was used to correct for multiple comparisons (Benjamini and Hochberg, 1995; Benjamini and Yekutieli, 2001). Differences were considered significant at false discovery rate adjusted p_fdr_<0.05.

For each brain region, we also calculated the number of voxels with significant β magnitude changes that survived at cluster-corrected threshold p<0.05 in individual subject analysis. The number of voxels with significant β magnitude changes were then normalized to the total number of voxels in each brain region. The percentage of voxels in each brain region with significant β magnitude changes, namely voxelβ_bER_, voxelβ_ΔPO2_ and voxelβ_ΔPCO2_, for RGE metrics bER, ΔPO_2_ and ΔPCO_2_ were calculated respectively. Individual subject brain volumes with regional voxelβ due to bER (voxelβ_bER_), ΔPO_2_ (voxelβ_ΔPO2_) and ΔPCO_2_ (voxelβ_ΔPCO2_) were obtained. One-sample t-tests were used to analyze regional voxelβ_bER_, voxelβ_ΔPO2_ and voxelβ_ΔPCO2_ in subject group. False discovery rate was used to correct for multiple comparisons (Benjamini and Hochberg, 1995; Benjamini and Yekutieli, 2001). Differences were considered to be significant at p_fdr_<0.05.

#### Group maps of regional ***Δ***BOLD associated with RGE Metrics vs. group functional connectivity maps with the seed at left precuneus

Seed-based analysis with the seed at the left precuneus was used to generate the functional connectivity map for individual subject. For each subject, after preprocessing of BOLD data, a low pass filtering at 0.03Hz was applied onto the brain volume with time courses of ΔBOLD. The time courses of ΔBOLD in the voxels within left precuneus were averaged. The averaged ΔBOLD time course of left precuneus was correlated with the ΔBOLD time course of each voxel in the brain volume. Pearson’s correlation coefficient was used to indicate the strength of correlation and converted to z scores using Fisher Z transformation. For the comparison with the regional β and voxelβ maps, the Fisher’s z scores in each of the 160 brain regions were averaged for each subject. Individual subject brain volumes with regional Fisher’s z scores from all the subjects who participated in MRI sessions were subjected to group analysis using one-sample t-test. False discovery rate was again used to correct for multiple comparisons (Benjamini and Hochberg, 1995; Benjamini and Yekutieli, 2001; Gold et al., 1998). Differences were considered to be significant at p_fdr_<0.05.

#### Dynamic analysis of coherence between bER and **Δ**BOLD in brain regions within DMN

To further verify the dynamic coupling between bER and CHF in DMN, we used the WTC method to examine the dynamic changes in coupling between the time courses of RGE metrics (bER, ΔPCO_2_, and ΔPO_2_) and ΔBOLD in three brain regions within DMN. They included inferior parietal lobule (IPL), posterior cingulate (PCC) and precuneus (PCun). The analysis procedures were the same as those described for the dynamic analysis of coherence between ΔCBFv and RGE metrics. The mean time-averaged coherence between each RGE metric and ΔBOLD from each of the three brain regions at the phase lag of 0±π/2 from all the subjects who participated in MRI sessions were averaged. Similarly, the mean time-averaged coherence between each RGE metric and ΔBOLD from each of the three brain regions at the phase lags of π±π/2 from the same group of the subjects were averaged.

#### Dynamic analysis of coherence between respiratory volume per unit time (RVT) and **Δ**BOLD in brain regions within DMN

To confirm that the respiratory-variation-related fluctuations reported by Birn et al. (2006) had little contribution to CHF within the DMN, we used the same WTC method to examine the dynamic changes in coupling between the time courses of RVT (respiration volume per time) and ΔBOLD in three brain regions within DMN (IPL, PCC and PCun). Ten out of 20 subjects RVT measurements with a respiratory bellow. According to the method described by Birn et al. (2006), RVT was computed by the difference between the maximum and minimum bellow positions at the peaks of inspiration and expiration respectively and this difference was then divided by the period of the respiratory cycle. The mean time-averaged coherence between RVT and ΔBOLD from each of the three brain regions at the phase lag of 0±π/2 from 10 subjects were averaged. Similarly, the mean time-averaged coherence between RVT and ΔBOLD from each of the three brain regions at the phase lag of π±π/2 from the same subjects were averaged.

#### Regional CVR quantification under exogenous CO_2_ challenge

Linear regression analysis was used to derive CVR from the time series of ΔBOLD and vasoactive stimulus when the subject was under exogenous CO_2_ challenge. We used the time course of P_ET_CO_2_ a regressor in a linear regression analysis. CVR was defined as the percent BOLD signal changes per mmHg change of P_ET_CO_2_. Therefore CVR was quantified by the coefficient of regression, i.e. the slope.

For each subject who participated in exogenous CO_2_ MRI scanning, CVR values derived from regressing ΔBOLD on P_ET_CO_2_ (CVR_CO2-PETCO2_) were separately averaged in each of the 160 brain regions parcellated by the software FreeSurfer. To study the CVR changes in group, one-sample t-tests were applied onto the brain volumes with regional CVR_CO2-PETCO2_. Differences were considered significant at false discovery rate adjusted p_fdr_<0.05.

## RESULTS

Subject demographics are shown in Table 1. The mean values and standard deviation of the time courses of P_ET_CO_2_, P_ET_O_2_, ΔPCO_2_, ΔPO_2_ and bER measured in the TCD and MRI sessions are summarized in Table 2.

**Table 2.** Mean values (SD) of P_ET_CO_2_, P_ET_O_2_, ΔPCO_2_, ΔPO_2_ and bER for all subjects who participated in the TCD sessions (n=13) (*left*), and for those who participated in the MRI sessions (n=20) (*right*).

| Subject<br>s | TCD |  |  |  |  | BOLD |  |  |  |  |
| --- | --- | --- | --- | --- | --- | --- | --- | --- | --- | --- |
|  | P<sub>ET</sub>CO<sub>2</sub><br>(mmHg) | P<sub>ET</sub>O<sub>2</sub><br>(mmHg) | ΔPCO<sub>2</sub><br>(mmHg) | ΔPO<sub>2</sub><br>(mmHg) | bER | P<sub>ET</sub>CO<sub>2</sub><br>(mmHg) | P<sub>ET</sub>O<sub>2</sub><br>(mmHg) | ΔPCO<sub>2</sub><br>(mmHg) | ΔPO<sub>2</sub><br>(mmHg) | bER |
| s1 | --- | --- | --- | --- | --- | 42.6 (2.1) | 99.4 (5.0) | 42.3 (2.3) | 52.6 (6.2) | 1.2 (0.1) |
| s2 | --- | --- | --- | --- | --- | 38.3 (1.3) | 111.5 (1.0) | 36.5 (2.0) | 35.6 (1.4) | 1.0 (0.1) |
| s3 | --- | --- | --- | --- | --- | 35.7 (3.8) | 109.0 (3.6) | 35.4 (4.4) | 41.7 (4.3) | 1.2 (0.2) |
| s4 | --- | --- | --- | --- | --- | 35.4 (1.2) | 114.5 (2.1) | 33.0 (1.2) | 37.1 (2.6) | 1.1 (0.1) |
| s5 | --- | --- | --- | --- | --- | 45.7 (2.1) | 113.9 (3.4) | 42.1 (3.2) | 29.4 (3.8) | 0.7 (0.1) |
| s6 | --- | --- | --- | --- | --- | 33.8 (4.0) | 118.8 (4.8) | 30.5 (4.0) | 25.1 (5.3) | 0.8 (0.1) |
| s7 | --- | --- | --- | --- | --- | 38.9 (1.8) | 116.2 (4.6) | 33.5 (1.7) | 30.8 (5.3) | 0.9 (0.1) |
| s8 | --- | --- | --- | --- | --- | 29.0 (1.8) | 103.9 (4.3) | 27.7 (1.9) | 50.9 (5.2) | 1.8 (0.1) |
| s9 | --- | --- | --- | --- | --- | 38.3 (1.3) | 98.0 (2.5) | 38.1 (1.5) | 55.3 (3.0) | 1.5 (0.1) |
| s10 | 39.1 (1.5) | 107.3 (3.2) | 39.4 (1.5) | 44.3 (3.7) | 1.1 (0.1) | 41.4 (1.8) | 103.8 (3.2) | 40.1 (1.8) | 43.0 (3.8) | 1.1 (0.1) |
| s11 | 39.1 (2.7) | 113.9 (4.1) | 38.5 (2.8) | 44.1 (4.6) | 1.1 (0.1) | 33.0 (2.9) | 111.1 (5.3) | 32.3 (3.0) | 37.1 (6.1) | 1.1 (0.1) |
| s12 | 30.6 (3.0) | 119.4 (6.2) | 29.5 (3.0) | 30.8 (7.0) | 1.0 (0.1) | 38.9 (2.5) | 107.8 (4.0) | 37.8 (2.6) | 39.6 (4.6) | 1.0 (0.1) |
| s13 | 37.2 (0.8) | 107.3 (2.0) | 37.3 (0.9) | 43.1 (2.5) | 1.2 (0.1) | 40.1 (0.9) | 108.1 (3.5) | 38.5 (1.0) | 41.9 (4.3) | 1.1 (0.1) |
| s14 | 39.4 (1.5) | 100.9 (3.2) | 38.3 (1.8) | 49.7 (4.0) | 1.3 (0.1) | 36.6 (0.7) | 108.7 (1.4) | 36.0 (0.8) | 40.0 (1.8) | 1.1 (0.0) |
| s15 | 34.7 (2.8) | 106.4 (4.1) | 34.9 (2.9) | 45.8 (4.6) | 1.3 (0.1) | 36.9 (1.9) | 114.4 (2.5) | 35.6 (1.9) | 34.9 (2.7) | 1.0 (0.0) |
| s16 | 25.9 (3.2) | 123.8 (4.5) | 26.0 (3.3) | 29.3 (5.2) | 1.1 (0.1) | 34.6 (1.3) | 117.8 (2.6) | 32.4 (1.4) | 34.5 (3.1) | 1.1 (0.1) |
| s17 | 35.9 (0.5) | 109.1 (1.2) | 35.3 (0.5) | 41.5 (1.4) | 1.2 (0.0) | 35.0 (0.6) | 112.2 (1.6) | 34.1 (0.7) | 38.8 (1.9) | 1.1 (0.0) |
| s18 | 34.0 (1.4) | 117.1 (3.9) | 33.4 (1.4) | 33.4 (4.6) | 1.0 (0.1) | 35.4 (3.2) | 115.3 (10.8) | 33.8 (3.3) | 38.5 (12.9) | 1.1 (0.3) |
| s19 | 33.9 (0.8) | 118.2 (2.2) | 29.3 (1.0) | 34.6 (2.9) | 1.2 (0.1) | 36.7 (1.0) | 113.5 (2.2) | 32.2 (1.1) | 39.2 (2.4) | 1.2 (0.1) |
| s20 | 32.4 (2.1) | 116.6 (3.1) | 29.1 (2.6) | 35.5 (4.0) | 1.2 (0.1) | 34.2 (1.9) | 114.4 (4.0) | 30.2 (2.1) | 38.6 (4.8) | 1.3 (0.1) |
| s21 | 36.1 (4.5) | 126.4 (4.9) | 28.9 (6.8) | 21.6 (6.5) | 0.7 (0.1) | --- | --- | --- | --- | --- |
| s22 | 32.4 (1.7) | 117.5 (2.8) | 28.1 (2.1) | 34.1 (3.5) | 1.2 (0.1) | --- | --- | --- | --- | --- |

### Correlation among bER, **Δ**PO_2_ and **Δ**PCO_2_

The correlations among the RGE metrics (bER, ΔPO_2_ and ΔPCO_2_) in TCD and MRI sessions are shown in Figure 1 and summarized in Table S1 (Supplementary materials). Correlation coefficients between ΔPO_2_ and ΔPCO_2_ ranged from 0.7 to 0.9 in TCD sessions and from 0.2 to 0.9 in MRI sessions, demonstrating that ΔPO_2_ and ΔPCO_2_ are not necessarily redundant in either TCD or MRI sessions. The range of correlation strength between ΔPO_2_ and ΔPCO_2_ in the MRI sessions was larger than that in TCD sessions. Correlation between bER and ΔPO_2_ was stronger than that between bER and ΔPCO_2_, suggesting bER is more driven by ΔPO_2_.

**Figure 1.**
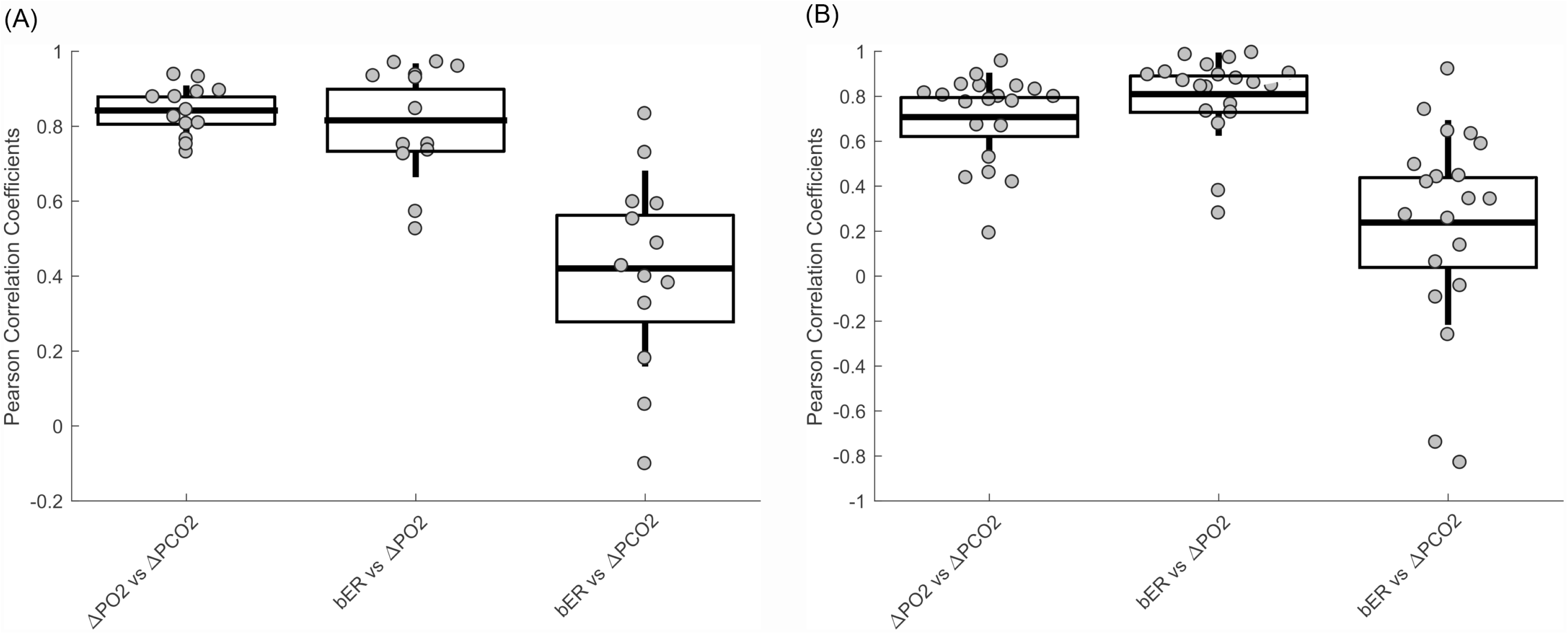
Correlations among breath-by breath RGE matrices (bER, ΔPO_2_ and ΔPCO_2_) in all subjects (A) who participated in TCD sessions (n=13), and (B) those who participated in MRI sessions (n=20). The major difference in the measurement of RGE metrics between TCD and MRI sessions was the posture of the subjects. In TCD sessions, subjects were in upright seated position, while they were in supine position in MRI sessions. Each gray circle represents the Pearson’s correlation coefficient from the correlation analysis of the time series of parameter pair shown on x-axis for each subject. The thick middle horizontal line, the box and the vertical rod represent the mean, standard deviation and 95% confidence interval of the group data respectively. The correlation coefficients from ΔPO_2_ vs ΔPCO_2_ varied from 0.7 to 0.9 in TCD sessions and from 0.2 to 0.9 in MRI sessions, indicating that ΔPO_2_ and ΔPCO_2_ are not necessarily redundant. The time series of bER had stronger correlation with ΔPO_2_ than that with ΔPCO_2_, although both ΔPO_2_ and ΔPCO_2_ contributed to changes of bER.

### Part 1

#### Correlation between **Δ**CBFv and RGE metrics

The time courses of ΔCBFv in right MCA of a representative subject during spontaneous breathing with and without applying low pass filtering at 0.03Hz are shown in Figure 2A. The time courses of bER, ΔPO_2_, ΔPCO_2_, P_ET_CO_2_ and P_ET_O_2_ acquired simultaneously with ΔCBFv for the same subject are shown in Figure 2B. All the time courses including CBFv, bER, ΔPO_2_, ΔPCO_2_, P_ET_CO_2_ and P_ET_O_2_ of a representative subject showed prominent oscillations with periods of 0.5 to 2 minutes which is equivalent to the frequency range of 0.008Hz to 0.03Hz. Oscillations of bER, ΔPO_2_, ΔPCO_2_, P_ET_CO_2_ were in phase with CBFv, while the oscillations of P_ET_O_2_ were out of phase. Oscillation amplitude of ΔPO_2_ could double that of ΔPCO_2_ (Figure 2A), where the standard deviation values indicating variations over the ΔPO_2_ time courses were larger than those over the ΔPCO_2_ time courses as shown in Table 2.

**Figure 2.**
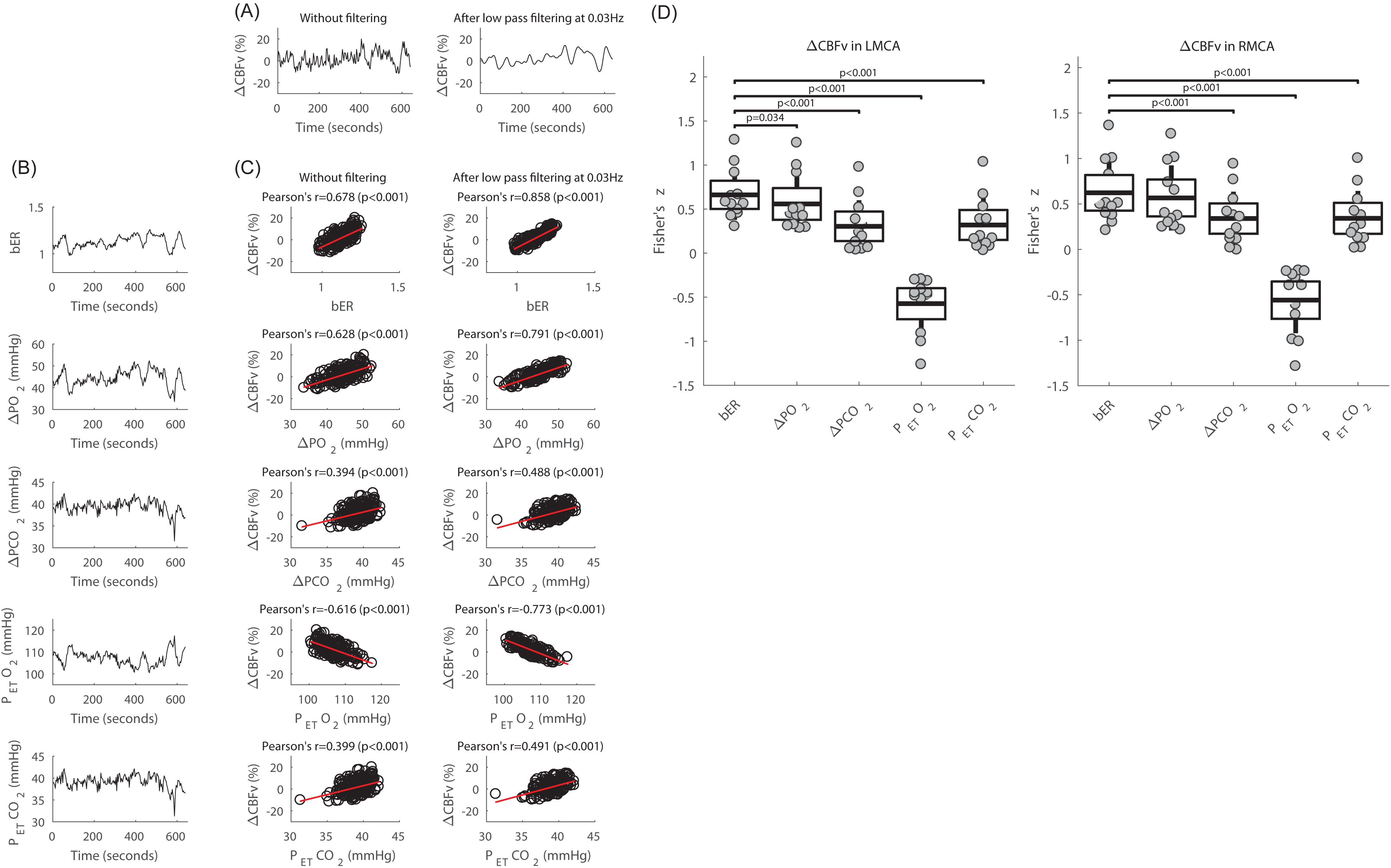
Correlation between the time courses of ΔCBFv in MCAs acquired with TCD and RGE metrics including bER, ΔPO_2_, ΔPCO_2_, P_ET_O_2_ and P_ET_CO_2_. (A) Time courses of Δ CBFv in right MCA of a representativo subject at rest, without filtering and after applying low pass filter at 0.03Hz. (B) Time courses of RGE metrics of bER, ΔPO_2_, ΔPCO_2_, P_ET_O_2_ and P_et_CO_2_ acquired simultaneously with ΔCBFv of the same subject in (A). (C) Correlation between ΔCBFv without filtering and the RGE metrics *(left),* and that between ΔCBFv with low pass filtering at 0.03Hz and RGE metrics *(right)* of the same subject in (A). (D) For all 13 subjects who participated in TCD sessions, paired comparisons of Fisher’s z scores between ΔCBFv in left MCA and bER, and those between ΔCBFv in left MCA and other RGE metrics besides bER *(left),* and the paired comparisons of Fisher’s z scores between ΔCBFv in right MCA and bER, and those between ΔCBFv in right MCA and other RGE metrics besides bER *(right).* Each gray circle represents the Fisher’s z score from the correlation analysis of ΔCBFv and the parameter shown on x-axis for each subject. The thick middle horizontal line, the box and the vertical rod represent the mean, standard deviation and 95% confidence interval of the group data respectively.

With the simple correlation analyses, Figure 2C shows that the correlation of bER with ΔCBFv was the strongest among all the RGE metrics. ΔPO_2_ followed relatively closely behind bER, and ΔPCO_2_ was the weakest in correlation with ΔCBFv. ΔPCO_2_ and P_ET_CO_2_ shared very similar correlation results with ΔCBFv as expected. The correlation coefficients between ΔCBFv and RGE metrics (bER, ΔPO_2_, ΔPCO_2_ and P_ET_CO_2_) for all 13 subjects who participated in TCD sessions are shown in Table S2 (Supplementary materials). The similar correlation of bER and ΔPO_2_ with ΔCBFv showed that ΔPO_2_ plays an important role in bER correlation with CHF. The correlation findings of RGE metrics with CHF extend and strengthen the correlation result among RGE metrics from Figure 1.

Applying a low pass filter of 0.03Hz to the time courses of ΔCBFv (Figure 2A, right panel) increased the strength of correlation of CBFv with bER, ΔPO_2_ and ΔPCO_2_ (Figure 2C, right panel) but did not change the order of which RGE metric was more correlated with ΔCBFv. Applying a low pass filter to CHF is a standard way to study resting state connectivity with BOLD data (Biswal et al., 1995).

In the group analysis, paired comparisons of Fisher z scores transformed from Pearson’s correlation coefficients between ΔCBFv and RGE metrics for all 13 subjects are shown in Figure 2D and Table S2 (Supplementary materials). The Fisher z scores of the correlation between ΔCBFv and bER were used as reference to compare with those of the correlation between ΔCBFv and other RGE metrics beside bER. ΔCBFv consistently showed stronger correlation with bER and ΔPO_2_ than with ΔPCO_2_ and P_ET_CO_2_. The differences revealed in paired comparisons again demonstrated that ΔPO_2_ and ΔPCO_2_ were not necessarily redundant.

#### Dynamic coherence between **Δ**CBFv and RGE metrics

The time-averaged coherence analyzed by WTC at phase lag of 0±π/2 which indicates positive correlation, and at phase lag of π±π/2 which indicates negative correlation, were plotted separately for all 13 subjects participated in TCD sessions (Figures 3). At the phase lag of 0±π/2, the mean time-averaged coherence between bER and ΔCBFv and that between ΔPO_2_ and ΔCBFv reached a value of 0.5 or above at the frequency range between 0.008 and 0.03Hz, while the mean time-averaged coherence between ΔPCO_2_ and ΔCBFv stayed below 0.1 (Figure 3A, top row). At the phase lag of π±π/2, the mean time-averaged coherence of all three RGE metrics with ΔCBFv stayed below 0.1 at the frequency range between 0.008 and 0.03Hz (Figure 3B, top row)).

**Figure 3.**
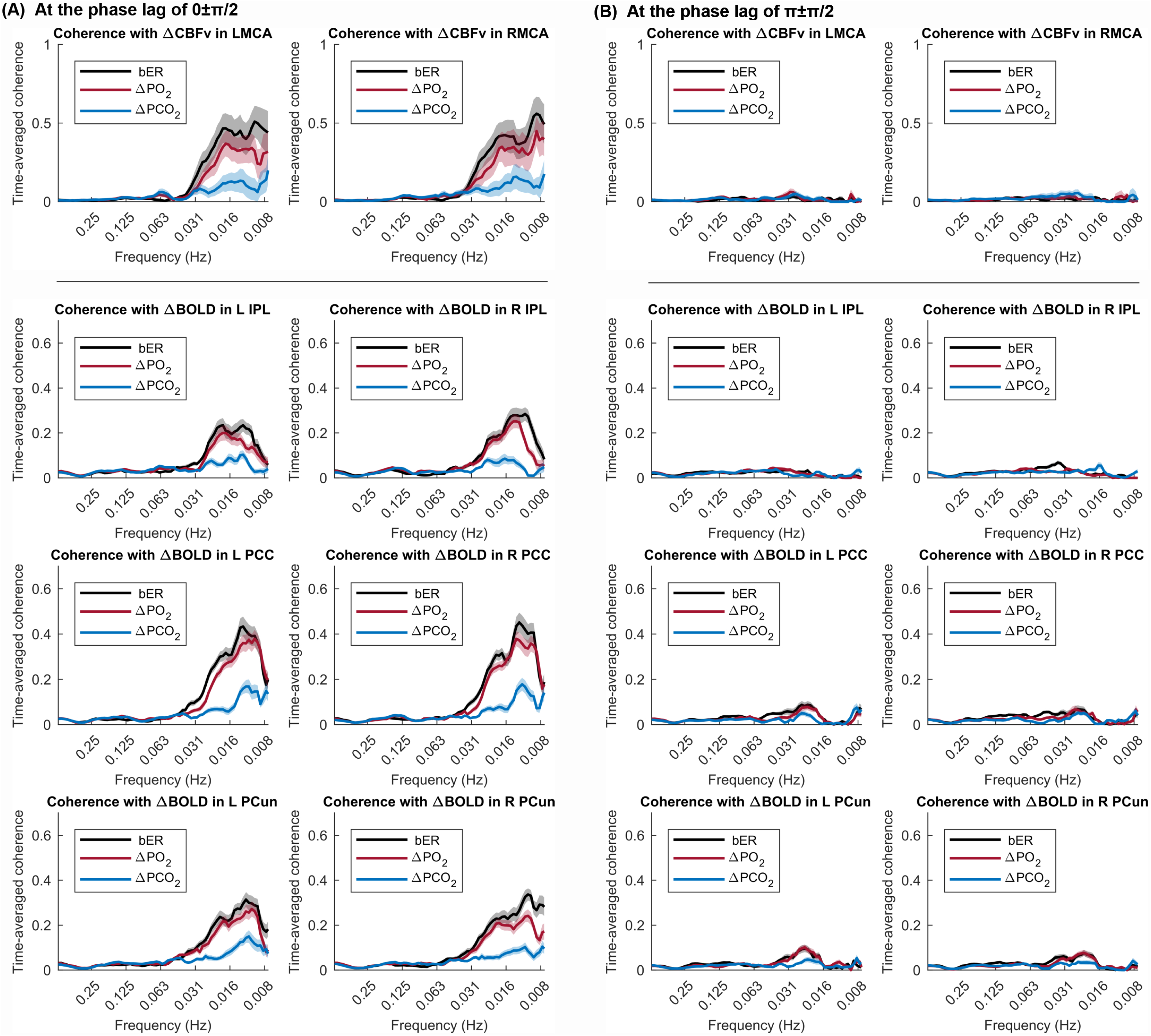
Distribution of time-averaged coherence between time series of RGE metrics (bER, ΔPO_2_ and ΔPCO_2_) and CHF (ΔCBFv and ΔBOLD) (A) at the phase lag of 0±π/2, and (B) at the phase lag of π±π/2. The mean time-averaged coherence in the frequency bandwidths from 0.008 to 0.25Hz were plotted (thick color lines). Color shaded areas represent standard error of the mean. Coherence between two time series at the phase lag of 0±π/2 indicates positive correlation, while negative correlation is represented by the coherence at the phase lag of π±π/2. Top panel shows the coherence with ΔCBFv in LMCA and RMCA in TCD sessions (n=13). The mean time-averaged coherence between bER and ΔCBFv reached 0.25-0.5 at the frequency range from 0.008Hz (1/128 seconds) to 0.03Hz (1/32 seconds), while the mean time-averaged coherence between ΔPCO_2_ and ΔCBFv stayed below 0.2. The lower panel shows the coherence between RGE metrics amd ΔBOLD in inferior parietal lobule (IPL), posterior cingulate (PCC) and precuneus (PCun) within default mode network (DMN) in MRI sessions (n=20). The mean time-averaged coherence between bER and ΔBOLD or between ΔPO_2_ and ΔBOLD was much higher than that between ΔPCO_2_ and ΔBOLD. In both TCD and MRI sessions, the coherence between RGE metrics and CHF (ΔCBFv and ΔBOLD) was minimal at the phase lag of π±π/2.

### Part 2

#### Association between RGE metrics and **Δ**BOLD in different brain regions

Group regional β magnitude changes were mapped for bER (β_bER_), ΔPO_2_ (β_ΔPO2_) and ΔPCO_2_ (β_ΔPCO2_) for all 20 subjects participated in MRI sessions (Figure 4A). ΔBOLD was found to be significantly associated with bER or ΔPO_2_ in most of the brain regions (Figure 4A) while association found between ΔBOLD and ΔPCO_2_ in a few brain regions (Figure S4 in Supplementary materials) did not survive after correcting for multiple comparisons (Figure 4A).

**Figure 4.**
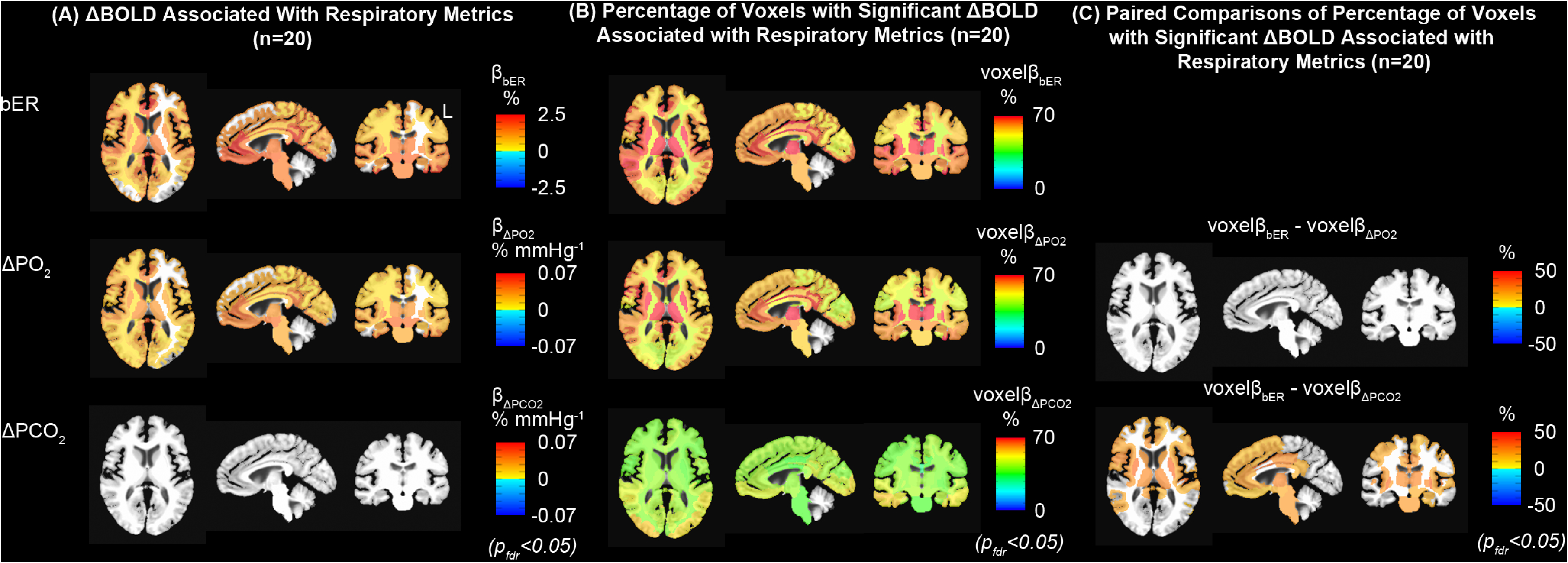
Regional association between ΔBOLD and RGE metrics for all subjects Included in the MRI sessions (n=20). False discovery rate was used to correct for multiple comparisons and significant association/differences were considered at false discovery rate adjusted p_fdr_<0.05. (A) Group maps showing regional ΔBOLD per unit change of bER, ΔPO2 and ΔPCO2 (β_bER_, β_ΔPO2_ and β_ΔPC02_). While significant changes of β_bER_ and β_ΔPO2_ (β_bER_, β_ΔPO2_ and β_ΔPCO2_) were shown in most of the brain regions, no significant changes of β_ΔPCO2_ were found. (B) Group maps showing regional percentage of voxels with significant changes of pbER, β_bER_,β_ΔPO2_ and β_ΔPCO2_ (voxelβ_bER,_ voxelβ_ΔPO2_ and voxelβ_ΔPCO2_). The changes of voxelβ_bER_, voxelβ_ΔPO2_ and voxelβ_ΔPCO2_were found in a decreasing order of voxelβ_bER_, voxellβ_ΔPO2_ and voxelβ_ΔPCO2_. (C) Paired comparisons of voxelβ maps showed that maps of voxelβ_bER_ at rest were not significantly different from those of voxelβ_ΔPO2_. However, brain regions including insula, caudate nucleus, thalamus, brainstem and medial frontal areas showed significant differences between voxelβ_bER_ and voxelβ_ΔPCO2_.

In addition to using the β magnitude, brain volumes with regional percentage of voxels having significant β changes for the regressors bER (voxelβ_bER_), ΔPO_2_ (voxelβ_ΔPO2_) and ΔPCO_2_ (voxelβ_ΔPCO2_) were analyzed in the same group of 20 subjects. Figure 4B shows that more than 60% voxels of gray matter regions showed significant association of ΔBOLD with bER and ΔPO_2_, while the percentage of significant voxels was smaller in white matter regions. A smaller percentage of gray and white matter voxels showed significant association of ΔBOLD with ΔPCO_2_. The maps of paired comparison between voxelβ_bER_ and voxelβ_ΔPO2_, showed no difference in percentage of significant voxels (Figure 4C) but those between voxelβ_bER_ and voxelβ_ΔPCO2_ showed a significant difference in percentage of significant voxels in a number of areas including subcortical brain regions, insula and brainstem. This again confirms the similarity between bER and ΔPO_2_ in their interaction with CHF.

#### Regional association between bER and **Δ**BOLD overlapped with many areas of the default mode network

Group statistical parametric maps of regional association between ΔBOLD and RGE metrics in gray matter and brainstem (Figure 5A) were used to compare with the group connectivity map with the seed at left precuneus (Figure 5B) for the same group of 20 subjects participated in MRI sessions. Brain regions showing significant association of ΔBOLD with bER and ΔPO_2_ including precuneus, posterior cingulate, anterior insula, caudate nucleus, superior temporal and inferior parietal regions overlapped with those in DMN reported by Raichle et al. (2001) as well as those reported by Yeo et al. (2011) and Choi et al. (2012). Additional regions such as posterior insula, putamen and occipital regions found in our group statistical parametric map were also shown in the findings by Raichle et al. (2001).

**Figure 5.**
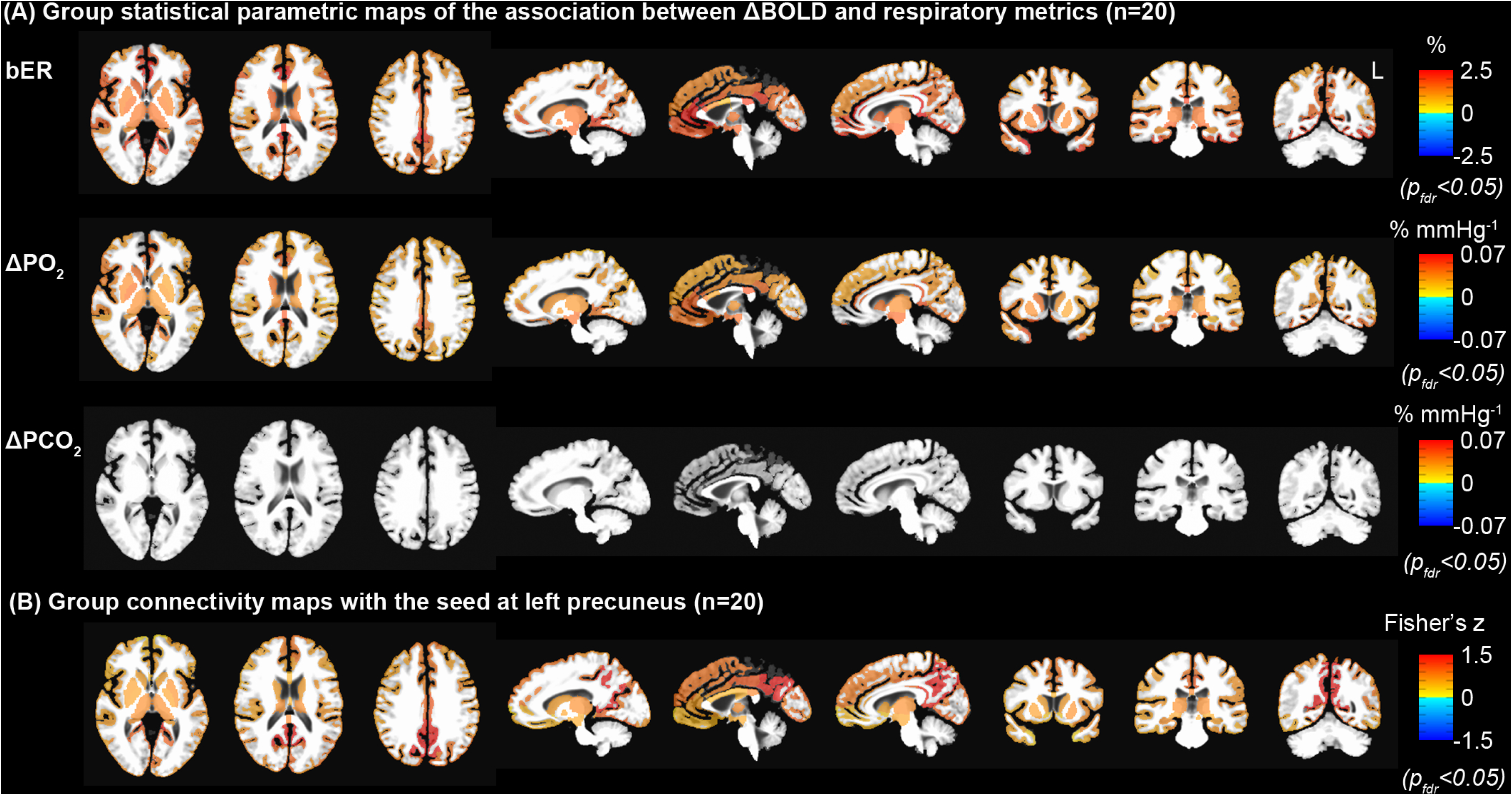
Comparison of statistical parametric maps derived from regression and connectivity analyses (n=20). False discovery rate was used to correct for multiple comparisons and significant associations/differences were considered at false discovery rate adjusted p_fdr_<0.05. (A) Group statistical parametric maps of the association between ΔBOLD and RGE metrics (bER, ΔPO_2_ and ΔPCO_2_) (n=20). The white matter was excluded for the comparison with the group connectivity map. (B) Group connectivity map with the seed at left precuneus (n=20). ΔPCO_2_ was demonstrated to be much weaker than bER and ΔPO_2_ in its association with regions of DMN. Comparing the top and bottom rows, similar regions of DMN were outlined with two independent methods of bER-CHF coupling and seed-based connectivity analysis.

Group analysis of individual connectivity maps demonstrated significant increased connectivity in brain regions similar to regions showing significant association between ΔBOLD and bER (Figure 5). The close match between Figures 5A and 5B from the same BOLD datasets but acquired with independent methods supports that bER is capable to outline the connectivity among brain regions in the DMN.

#### Dynamic coherence between bER and **Δ**BOLD in brain regions within DMN

We used WTC analysis to verify the dynamic coupling between bER and BOLD signal fluctuations in DMN regions such as left and right inferior parietal lobule (IPL), left and right posterior cingulate (PCC), and left and right precuneus (PCun) in all 20 subjects participated in MRI sessions. At the phase lag of 0±π/2, we showed that the mean time-averaged coherence between bER and ΔBOLD and that between ΔPO_2_ and ΔBOLD reached a value of 0.3 or above in general (Figure 3A, lower panel) at the frequency range between 0.008 and 0.03Hz, while the mean time-averaged coherence between ΔPCO_2_ and ΔBOLD peaked around 0.2 or below (Figure 3A, lower panel). At the phase lag of π±π/2, the mean time-averaged coherence of all three RGE metrics with ΔBOLD stayed below 0.1 at the frequency range between 0.008 and 0.03Hz (Figure 3B, lower panel).

#### Dynamic coherence between RVT and **Δ**BOLD in brain regions within DMN

To confirm that the RVT had little contribution to CHF within the DMN, we used the WTC method to examine the dynamic coupling between RVT and ΔBOLD extracted from IPL, PCC and PCun in 10 subjects who had their respiratory tracings measured by a respiratory bellow. We showed that the mean time-averaged coherence between RVT and ΔBOLD at the phase lag of 0±π/2 was increased between 0.016 and 0.031Hz, and reduced at the frequency below 0.016Hz where the mean time-averaged coherence started to increase at the phase lag of π ±π/2 (Figure S3 in Supplementary materials). The frequency distribution of coherence of CHF with RVT is shown to be very different from that with bER.

#### Regional CVR maps under exogenous CO_2_ challenge and during spontaneous breathing at rest

Under exogenous CO_2_ challenge, most of the brain regions showed increased CVR_CO2-PETCO2_ in the subject group especially thalamus, insula and putamen (Figure S5 in Supplementary materials). During spontaneous breathing, increased β_bER_ was found in most of the brain regions, while no significant changes of β_ΔPCO2_ were shown in most of the brain regions. The CVR map during spontaneous breathing indicated by β_bER_ changes resembled the CVR map under exogenous CO_2_ challenge indicated by CVR_CO2-PETCO2_.

## DISCUSSION

While there has been no previous study on the dynamic interaction between low frequency fluctuations of resting state CHF and those of all three RGE metrics of bER, ΔPO_2_ and ΔPCO_2_, results in this study support our hypothesis for such an interaction. Independent of the capability of our current study to investigate any specific underlying mechanism for such an interaction, we uncovered the dynamic brain-body interaction between resting state CHF and our designated RGE metrics, with CHF showing much stronger interaction with bER and ΔPO_2_ than with ΔPCO_2_ in both the time and frequency domains. The dynamic changes in bER, ΔPO_2_ and ΔPCO_2_ were characterized by prominent oscillations with periods of 0.5 to 2 minutes, consistent with the coherence found between RGE metrics and CBFv in the frequency range of 0.008-0.03Hz. In the time-frequency wavelet transform coherence analysis, the coherences of CHF with bER and of CHF with ΔPO_2_ were much stronger than the coherence of CHF with ΔPCO_2_ in the frequency range between 0.008Hz and 0.03Hz. Brain regions with the strongest bER-CHF coupling overlapped with many areas of DMN. Taken together, our dynamic brain-body interaction provides evidence that fluctuations related to RGE metrics are associated with resting state CHF at low frequency. No matter whether the contribution of bER fluctuations to resting state CHF is considered as ‘physiological noise’ similar to respiratory variability (Birn et al., 2006) and heart rate variability (Chang et al., 2013) to be removed, or as ‘useful signals’ for the evaluation of cerebrovascular reactivity (CVR) to the changes of systemic/endogenous gases during spontaneous breathing (Golestani et al., 2016; Liu et al., 2017b), bER may offer a better choice of regressor than ΔPCO_2_.

### Oscillations of **Δ**PO_2_ and **Δ**PCO_2_ are interdependent but not necessarily redundant

ΔPO_2_ and ΔPCO_2_ were often considered to be redundant (Chang and Glover, 2009). In the present study, the average value of the correlation strength of around 0.8 and the relatively large spread of the correlation between ΔPO_2_ and ΔPCO_2_ demonstrated that while ΔPO_2_ and ΔPCO_2_ were interdependent, they were not necessarily redundant to each other (Figure 1). This is consistent with the findings on healthy infants reported by Cernelc et al. (2002) that different long-range correlations were found between time series of P_ET_CO_2_ and P_ET_O_2_. Respiratory data acquired on 12 subjects in the study by Lenfant (1967) were consistent with ours showing that end-tidal partial pressures of O_2_ and CO_2_ were correlated but not redundant. Our bER was also shown to have stronger correlation with ΔPO_2_ than with ΔPCO_2_, suggesting that bER is more driven by ΔPO_2_. The spread of correlation strengths between ΔPO_2_ and ΔPCO_2_ as well as those between bER and ΔPO_2_ were different between TCD and MRI sessions. It is possible that correlation strengths changed in response to different environments which could include the postures of sitting up vs. lying down, the amount of noises, or the level of stress from the surroundings. Participants were sitting in an open and quiet environment in the TCD sessions while they were lying down in supine position in a noisy environment of the MRI scanner bore. Many previous studies reported that a change from the supine posture to sitting upright was associated with a redistribution of both blood flow and ventilation in the lungs, which affected the arterial PO_2_ (Amis et al., 1984; Glaister, 1967; Petersson et al., 2009; Remolina et al., 1981). Our claim that the extent of non-redundancy can vary with posture reinforce only the notion that there are many different ways for ΔPO_2_ and ΔPCO_2_ to become non-redundant. While it is interesting to observe a difference in the interaction between individuals and the environment, the details of such mechanisms are outside the scope of our current study.

### Dynamic coupling between CHF and RGE metrics of bER, **Δ**PO_2_ and **Δ**PCO_2_

Correlation and coherence analyses showed the dynamic coupling between CHF (ΔCBFv and ΔBOLD) and RGE metrics of bER, ΔPO_2_ and ΔPCO_2_ in both TCD (Figures 2 and 3) and MRI sessions (Figures 3 and 4). Had ΔPO_2_ and ΔPCO_2_ been redundant, their correlation and coherence with CHF would be expected to be the same, and there should be weak correlation and coherence of CHF with bER. Our results differed from such expectations. Comparing with ΔPO_2_ and bER, ΔPCO_2_ had moderate to weak correlation with ΔCBFv. Only a few brain regions in caudate nuclei, right frontal and temporal areas showed association between ΔPCO_2_ and ΔBOLD (Figure S4 in Supplementary materials) but such association did not survive after correcting for multiple comparisons (Figure 4). We also found larger oscillation amplitude of ΔPO_2_ in comparison with that of ΔPCO_2_ (Figure 2 and Table 2). This finding is consistent with that in the study by Lenfant (1967) which reported that the oscillation amplitude of end-tidal O_2_ time course was larger than that of end-tidal CO_2_ time course in healthy adults. The interdependent but non-redundant oscillations of ΔPO_2_ and ΔPCO_2_ during spontaneous breathing at normoxic level are supported by the findings of feedback loops involving the interaction among chemoreceptors and blood gases demonstrated in many earlier studies on animal models (Biscoe et al., 1970; Honda et al., 1963; Kumar and Prabhakar, 2012; Lahiri and DeLaney, 1975a; Lahiri et al., 1978). While most of these studies on the role of PO_2_ to stimulate peripheral chemoreceptors at the carotid body had been focused on hypoxia at PO_2_ level less than 60 mmHg (Kumar and Prabhakar, 2012; Weir et al., 2005) where chemoreceptor activities rose quickly in a hyperbolic fashion, Biscoe et al. (1970) showed that peripheral chemoreceptor activities were present from normoxia to hyperoxia up to arterial PO_2_ level of 190 mmHg and beyond. Lahiri et al. (1978) reported that the stimulus thresholds of arterial PO_2_ and PCO_2_ for peripheral chemoreceptors were largely interdependent under normoxic conditions where a drop in arterial PO_2_ was routinely accompanied by increased chemoreceptor activities as well as an enhanced sensitivity of carotid chemoreceptors to arterial PCO_2_. While research on the mechanisms of interaction between peripheral and central chemoreceptors to optimize systemic gases is on-going (Cummins et al., 2019; Kumar and Prabhakar, 2012; Rocha and Branco, 1998), our findings here suggest that PO_2_ and PCO_2_ work synergistically to regulate blood supply to the brain. bER is one of the appropriate metrics to be used to study such synergism of ΔPO_2_ and ΔPCO_2_. The superiority of bER over ΔPCO_2_ in correlating with ΔCBFv is likely to be attributed partly to the ratio format of bER enabling a reduction of ventilatory fluctuations and partly to the physiological role of bER which takes into account the interaction between ΔPO_2_ and ΔPCO_2_. On the other hand, while any interaction between ΔPCO_2_ and CHF would necessarily be accompanied by an interaction between ΔPO_2_ and CHF, the mechanisms behind the superiority of ΔPO_2_ over ΔPCO_2_ in association with CHF, especially at the lower end of the frequency range of 0.008-0.03Hz, remain unclear. In addition to our studying of healthy subjects, exploring the interaction between CHF and RGE metrics in patients with known disorders may provide opportunities to increase the understanding of the stronger coupling of CHF with bER and ΔPO_2_ than with ΔPCO_2_.

### Brain Regions with the Strongest bER-CHF Coupling Overlapped with Many Regions of Default Mode Network

We used the resting state connectivity method with the seed at the left precuneus to outline the areas within DMN (Figure 5B). We found that brain regions with the strongest association between bER and ΔBOLD (Figure 5A) overlapped with many areas of DMN. The strong association may be attributed to DMN showing higher metabolic and hemodynamic activities at rest than other parts of the cortex (Raichle et al., 2001). Stronger association of DMN activities with bER than with ΔPCO_2_ could be attributed to bER’s stronger association with CHF. Fluctuations of systemic gases can be under the influences of ambient (exogenous) gases and/or systemic (endogenous) gases due to metabolism as well as feedbacks between chemoreceptors. Since brain is one of the major shareholders of systemic blood flow (Markus, 2004) and metabolism (Kuzawa et al., 2014; Wang et al., 2010) at rest, any mechanisms that help to preserve cerebral metabolism and CBF within normal range in the homeostatic regulatory process (Modell et al., 2015) may leave a signature in the CHF.

In our resting state experiments, CHF were measured on participating individuals at rest during spontaneous breathing. No cognitive, motor, sensory, visual and gas challenges were applied to the participants. The internal environment of the body was steady and major body organs had constant autonomic communication with the brain back and forth. At rest, background cerebral neurovascular coupling activities are characterized by DMN (Raichle et al., 2001) and RGE metrics can be related to basal activities of the whole body. Hence it would be reasonable for RGE metrics to couple better with DMN than with other brain networks. bER which showed the strongest correlation with CHF is therefore expected to interact more with DMN. The bER-DMN coupling is consistent with findings in previous studies that CBF and DMN connectivity were altered in individuals with disrupted systemic metabolism (Chen et al., 2017; Ishibashi et al., 2018; Ishibashi et al., 2016; Li et al., 2017; Liu et al., 2017a; Liu et al., 2019; Lockwood et al., 1991; Qi et al., 2014).

### Physiological processeses which may be associated with the coherence between CHF and bER at the low frequency range of 0.008-0.03Hz

The CHF coherence with RGE metrics at the low frequency range of 0.008-0.03Hz may be associated with low frequency physiological processes in the brain that are grouped in B-wave frequency bandwidth. B-waves with a period of 0.5 to 2 minutes have been reported to be related to autoregulation of microvasculature, spontaneous rhythmic oscillations in intracranial pressure (ICP), and intrinsic brainstem rhythm that leads to cerebral blood volume modulation (Lundberg, 1960; Spiegelberg et al., 2016). B-waves may also independently reflect neurovascular coupling process by altering vascular diameters to ensure the delivery of O_2_ and other circulating metabolites through the contractile properties of pericytes or vascular smooth muscle cells (Hill et al., 2015; Mulligan and MacVicar, 2004; Takano et al., 2006; Zonta et al., 2003). Since oscillations of B-waves can be associated with or without the change of arterial CO_2_ (Spiegelberg et al., 2016). it would be interesting to pursue further the relationship between B-waves and all three of our RGE metrics as well as resting state CHF.

The coherence of CHF with RGE metrics at frequency of 0.008-0.03Hz may also be related to a proposed mechanism to remove cerebral metabolic waste. In the glymphatic model (Iliff et al., 2012), cerebral metabolic waste was hypothesized to be cleared from the brain via the perivascular space by vascular pulsation. The actual type of vascular pulsation remains to be determined. To describe possible mechanisms for glymphatic convection, Kiviniemi et al. (2016) identified a very slow pulsation found in cerebrospinal fluid (0.001-0.023Hz), which is again in the same frequency range of the oscillations of the RGE metrics.

In the peripheral circulation, laser Doppler measurements showed that the endothelial activity oscillated at the frequency of 0.0096-0.021Hz (Bernjak et al., 2008; Stefanovska et al., 1999), which is within the same low frequency range of CHF and of fluctuations of RGE metrics with a special interest on bER based on our results. Is there any direct relationship between oscillatory cycles of peripheral endothelial activity and CHF? Considering the model of diving reflex (Gooden, 1994) where CBF increase is associated with peripheral vasoconstriction, is it possible that the decrease in peripheral blood flow is mirrored by increase in CBF in a homeostatic process during spontaneous breathing at rest? Future research can be directed to explore the association between bER, CHF and peripheral (skin) blood flow oscillations at the low frequency range of 0.008–0.03 Hz.

Separate from B-waves and glymphatic model, we do not consider heart rate (HR) or heart rate variability (HRV) to be a major contributing factor for the role of RGE metrics in CHF. Even though HR and HRV had been reported to contribute to low frequency fluctuations, the peaks of coherence between ΔBOLD and HR/HRV were at the frequency of 0.05Hz or above (period ∼30-42 seconds) (Chang et al., 2013; Shmueli et al., 2007) which were different from our findings that RGE metrics were coherent with CHF between 0.008 and 0.03Hz.

### Ventilatory volume fluctuations are not the primary origin of our results

We should point out that ventilatory volume fluctuations are unlikely to be the cause for our observed results. Respiratory variability, namely changes in respiration volume per time or RVT, had been discussed as a possible source for DMN activities (Birn et al., 2006). RVT shows strong correlation with change in respiratory volume (Van den Aardweg and Karemaker, 2002). However, a number of considerations showed that RVT did not provide an answer to our findings on the discrepancy observed between ΔPO_2_ and ΔPCO_2_ in their interaction with CHF. First, RVT would be expected to have the same effect on the time courses of ΔPO_2_ and ΔPCO_2_. Secondly, bER, being a ratio of ΔPO_2_/ΔPCO_2_, reduces the contribution of fluctuations of ventilatory volumes to the interaction between bER and CHF. Previous studies did show that the relationship between the time courses of P_ET_CO_2_ and RVT was unclear (Golestani et al., 2015; Van den Aardweg and Karemaker, 2002). The authors in these studies showed that changes in RVT had weaker correlation with BOLD signal changes when compared with P_ET_CO_2_ (Golestani et al., 2015; Vogt et al., 2011). Such results are not surprising as P_ET_CO_2_ is supposed to be a more direct mechanism than ventilation to modulate CHF (Birn et al., 2006; Chang and Glover, 2009). Besides, the time course of RVT is less than ideal as the numerator of RVT acquired with a respiratory bellow indicates the chest excursion, while respiratory volume is the volume of air breathing in or out. Our coherence analysis showed that ΔBOLD at DMN regions were less coherent with RVT than with bER or ΔPO_2_ (Figure S3). RVT showed coherence with ΔBOLD at the phase lag of 0±π/2 between 0.016 and 0.031Hz but the coherence was between 0.008 and 0.016Hz at the phase lag of π±π/2. The coherence findings of RVT were also different from those of ΔPCO_2_. The frequency bandwidths and the phase differences seen in our coherence of RVT with ΔBOLD are consistent with those reported by Van den Aardweg (2002) even though his team was measuring respiratory volume from a facemask. While it is possible that coherence frequency bandwidths at different phase lags are referring to the responses from different chemoreceptors (e.g. central vs. peripheral), the relationship between the time courses of P_ET_CO_2_ and RVT is still unclear.

### Potential application of bER in the evaluation of CVR

While bER was prominently coupled with DMN, bER was also coupled, albeit to a less extent, with the rest of voxels throughout the whole brain. Therefore bER can be used instead of ΔPCO_2_ as a superior regressor to evaluate CVR to spontaneous breathing (Figure S5 in Supplementary materials). The question is whether the expected small perturbations provided by RGE metrics during spontaneous breathing could have any clinical utility. Previous study on the successful identification of local vascular deficits of moyamoya patients (Liu et al., 2017b) had already been reported for CVR to P_ET_CO_2_ obtained in spontaneous breathing. Future studies will clarify the sensitivity and benefits of imaging local vascular deficits using bER instead of P_ET_CO_2_ from spontaneous breathing.

Imaging of bER-CHF coupling may open up opportunities for potential clinical diagnosis and intervention. The clinical potential of RER at rest has been reported for decades. RER, the reciprocal of the steady state mean value of bER, has been reported to change significantly with age (Rizzo et al., 2005) and with diseases attributed to respiratory (Cai et al., 2003), liver (Lockwood et al., 1991), cardiac (Krauss and Auld, 1975) and neuronal (Dupuis et al., 2004) dysfunctions. Instead of focusing on the steady state results, our manuscript centers more on the dynamic fluctuations of bER, as a separate metric of interest. bER-DMN coupling may offer an alternative approach to map brain regions interacting with systemic homeostatic processes besides resting state connectivity analysis for the connections among brain regions (Greicius et al., 2003) and task-induced negative BOLD responses (Raichle et al., 2001). Since abnormal DMN can be an indicator of neuronal disorders (Greicius et al., 2004; Raichle, 2015), future studies on different patient populations may help clarify the meaning of abnormal bER-DMN coupling. Our brain-body coupling framework may apply in the investigation of brain responses to voluntary respiratory techniques like meditation, yoga breathing or tai-chi (Chaya et al., 2006; Garcia et al., 2018; Kesterson and Clinch, 1989; Ray et al., 2011; Sood et al., 2007; Tai et al., 2018; Wells et al., 2013; Zheng et al., 2015) to manipulate RGE for intervention purpose. This also includes the Bi-level Positive Airway Pressure (BiPAP) (Carlucci et al., 2015; Khamankar et al., 2018; Xiang et al., 2009) that applies to sleep apnea or Amyotrophic Lateral Sclerosis (ALS).

### Technical issues that have been addressed and will need to be addressed

In this study, in addition to BOLD-fMRI, we used TCD to acquire CHF. TCD has several advantages including its high temporal resolution with less concern on the aliasing effect of high frequency hemodynamic signal. Potential technical confounds like gas sampling at two gas analyzers and technical delay on the time courses were also addressed in the data acquisition protocol. In the gas sampling circuit, the same gas sample volume was used in both CO_2_ and O_2_ analyzers at the same gas sampling flow rate. The technical delay on the time courses of gas measurements due to transit time of respiratory gases and the response of the equipment were corrected in the preprocessing of the physiological signals. In the preprocessing of fMRI data, we did not apply RETROICOR (Glover et al., 2000) to reduce ventilatory and cardiac ‘interference’ in the BOLD signals because heart rate was reported to contribute to BOLD signals mainly at 0.3Hz or above (Shmueli et al., 2007) which was above the frequency bandwidths of interest in this study. The preprocessing step for BOLD signals like data smoothing (Figure 2) did not change the ranking of ΔBOLD with bER, ΔPO_2_ and ΔPCO_2_.

Even though ΔPO_2_ is related to the partial pressure of utilized O_2_ of the whole body, ΔPO_2_ and ΔPCO_2_ by themselves are not yet equal to O_2_ uptake (VO_2_) and CO_2_ release (VCO_2_). O_2_ uptake and CO_2_ release require extra elements like respiratory minute volume (sometimes normalized for body weight) and are normally evaluated as a steady state of RGE data obtained by averaging the breath-by-breath signal over many minutes. A measurement of breath-by-breath VO_2_ and VCO_2_ inside the MRI scanner in addition to collection of ΔPO_2_ and ΔPCO_2_ was not included in our study. Given the physical constraint of the MRI settings, we collected ΔPO_2_ and ΔPCO_2_ using a nasal tubing. We will leave the possibility of measuring VO_2_ and VCO_2_ during MRI to future neuroimaging research. However, under specific conditions, some information extracted from ΔPO_2_ and ΔPCO_2_ can be related to that from VO_2_ and VCO_2_. As mentioned above, bER has the explicit influence of ventilatory volume suppressed when we are taking the ratio of ΔPO_2_ and ΔPCO_2_ from the same individual. We therefore consider the possibility of bER or equivalently ΔPO_2_/ΔPCO_2_ being a surrogate for breath-by-breath VO_2_/VCO_2_ at rest.

As the vasodilatory role of CO_2_ to modulate CBF is an entrenched belief, it is natural to gravitate towards the explanation that association between RGE metrics and CHF must be resulting from ‘the effect of CO_2_ which reflects in O_2_’. However, the model of CO_2_ being the sole agent to interact with CHF remains to be reconciled with our findings of non-redundancy between ΔPO_2_ and ΔPCO_2_ in this study. Therefore, further investigations on the roles of bER, ΔPO_2_ and ΔPCO_2_ in their association with CHF would be warranted.

## CONCLUSION

Independent of our current knowledge to clarify the physiological mechanisms to explain the interaction between CHF and RGE metrics of bER, ΔPO_2_ and ΔPCO_2_, our findings provide evidence that fluctuations of RGE metrics are associated with resting state CHF at low frequency between 0.008Hz and 0.03 Hz, with bER being the superior RGE metric, followed by ΔPO_2_ and then by ΔPCO_2_. Brain regions with the strongest bER-CHF coupling overlap with many areas of DMN. In addition to offering a physiological model to characterize the contribution of gas exchange elements in the low frequency resting state fluctuations, our findings provide new directions to the study of brain-body interaction. Assessing the validity of the popular CO_2_-only hypothesis as well as other alternative hypotheses to explain our results of bER-CHF coupling remains an on-going project.

## ACKNOWLEDGMENTS

Our team appreciates the expertise physiologic advices provided by Dr. Robert Banzett.

## AUTHOR CONTRIBUTIONS

Data collection (STC, KCE, TYS, JS, KKK), data analysis (STC, KCE, TYS, YPZ, KKK), study design (STC, KCE, AVK, BRR, YPZ, KKK), manuscript preparation (STC, BRR, YPZ, KKK).

