## Supplemental Information for "Dynamic Brain-Body Coupling of Breath-by-Breath O_2_-CO_2_ Exchange Ratio with Resting State Cerebral Hemodynamic Fluctuations"

**Figure S1**

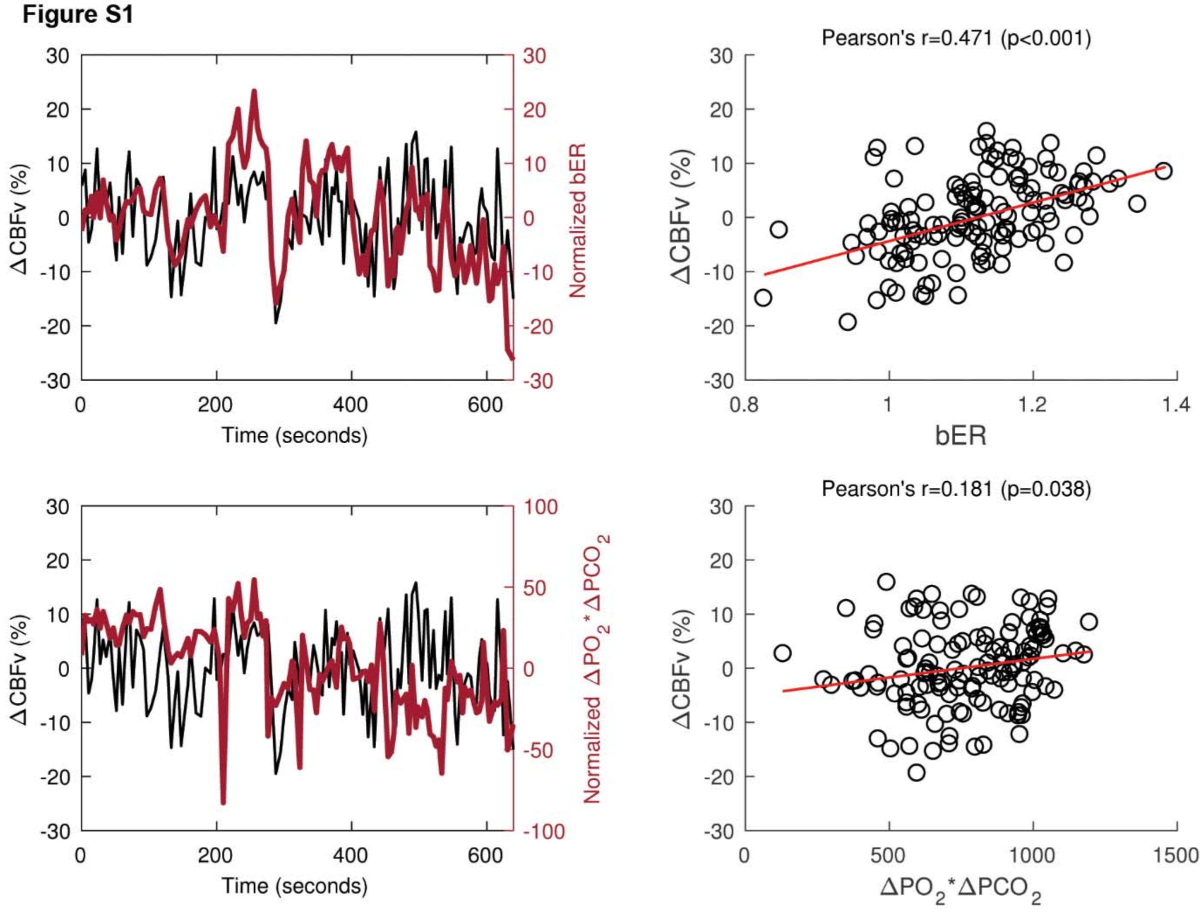


Figure S1. Correlation of ΔCBFv with the ratio as well as the product of $\Delta PO_2$ and $\Delta PCO_2$ in a representative subject. Time series of ΔCBFv and bER normalized to its mean (*upper left*). Moderate correlation was shown between ΔCBFv and bER (*upper right*). Time series of ΔCBFv and the product of $\Delta PO_2$ and $\Delta PCO_2$ ($\Delta PO_2$*$\Delta PCO_2$) normalized to its mean (*lower left*). Weak correlation was shown between ΔCBFv and $\Delta PO_2$*$\Delta PCO_2$ (*lower right*).

Figure S2

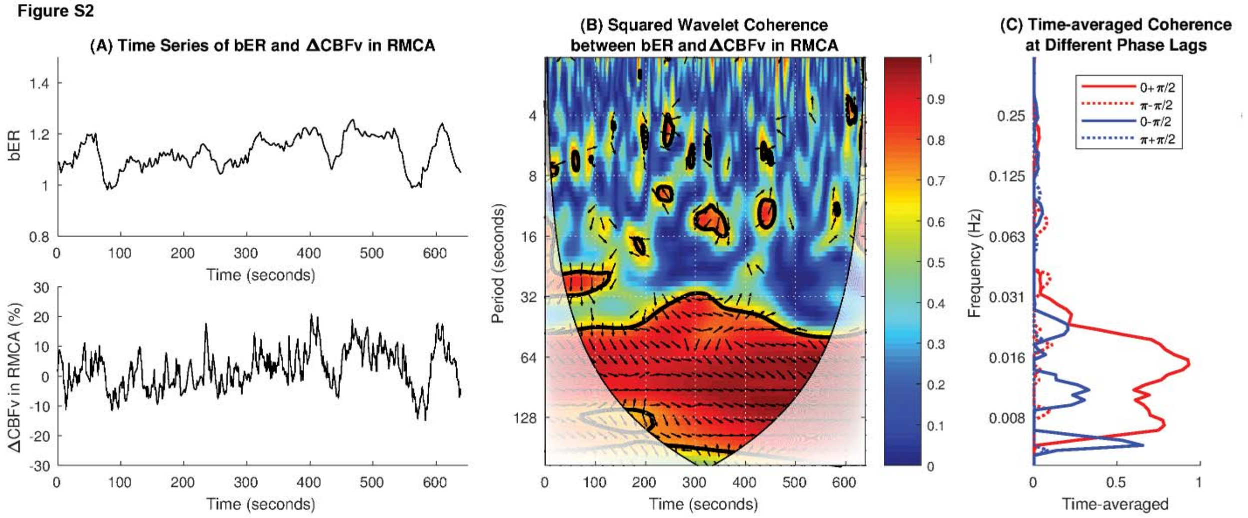


Figure S2. Example of wavelet transform coherence analysis. (A) Time series of bER and ΔCBFv measured in right MCA of a representative subject at rest. (B) The squared wavelet coherence between these two time series. Squared wavelet coherence is plotted with x-axis as time and y-axis as scale which has been converted to its equivalent Fourier period. The magnitude of wavelet transform coherence ranges between 0 and 1, where warmer color represents stronger coherence and cooler color represents weaker coherence. Areas inside the 'cone of influence', which are locations in the time-frequency plane where edge effects give rise to lower confidence in the computed values, are shown in faded color outside of the conical contour. The statistical significance level of the wavelet coherence is estimated using Monte Carlo methods and the 5% significance level against red noise is shown as thick contour. The phase angle between the two time series, with bER leading ΔCBFv, at particular samples of the time-frequency plane is indicated by an arrow (rightward pointing arrows indicate that the time series are in phase or positively correlation, leftward pointing arrows indicate anticorrelation and the downward pointing arrows indicate phase angles of π/2). There are four different ranges of phase lags: 0+π/2, π-π/2, 0-π/2 and π+π/2. (C) Time-averaged coherence at four different phase lags of 0+π/2, π-π/2, 0-π/2 and π+π/2. At each phase lag range, time-averaged coherence was defined as the total significant coherence at each scale where the wavelet coherence magnitude exceeded 95% significance level, normalized by the maximum possible coherence outside the cone of influence, i.e. inside the conical contour, at that particular scale.

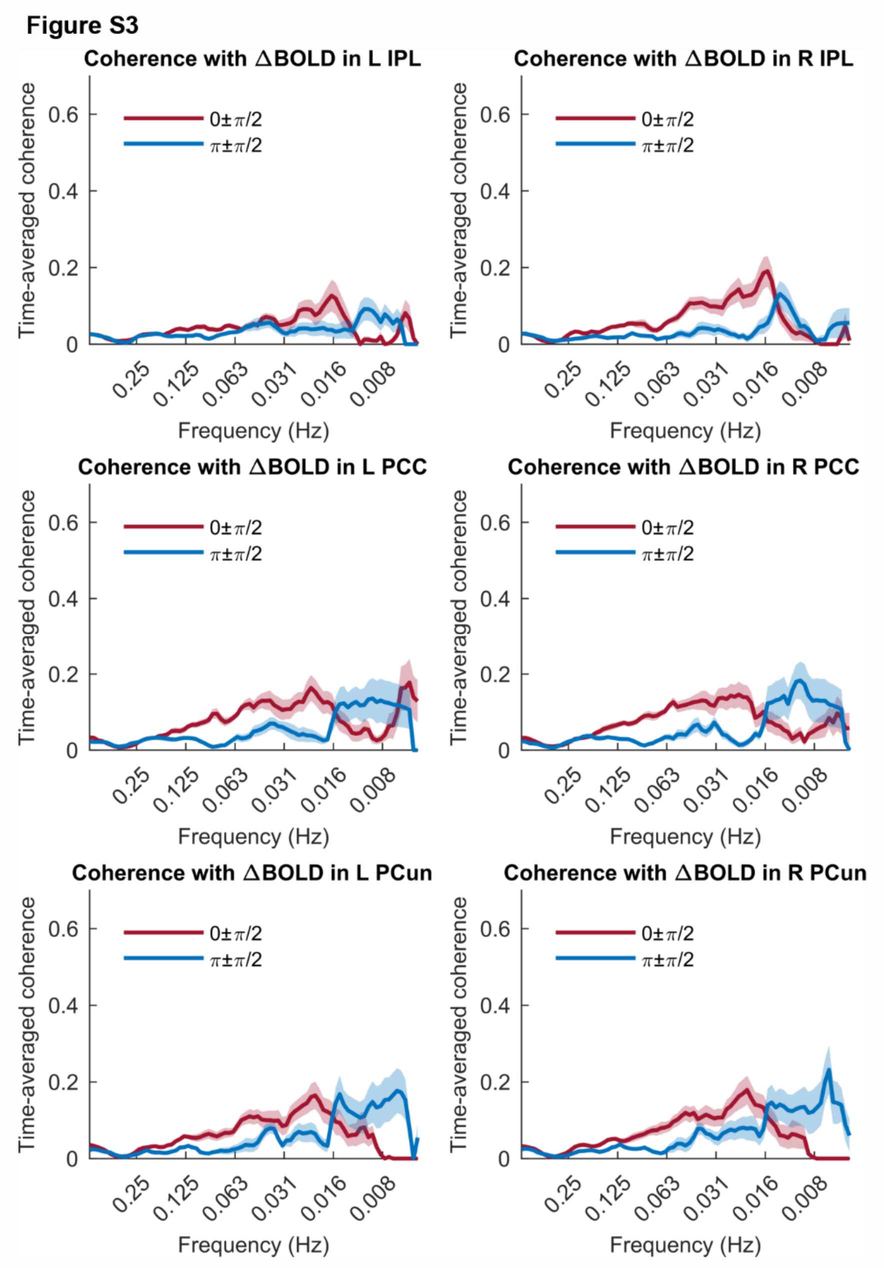


Figure S3. Distribution of time-averaged coherence between time series of respiratory volume per unit time (RVT) and ΔBOLD at the phase lags of 0±π/2 and π±π/2 in the inferior parietal lobule (IPL), posterior cingulate (PCC) and precuneus (PCun) of the left brain (*left panel*) and of the right brain (*right panel*) (n=10). The mean time-averaged coherence in the frequency bandwidths from 0.008 to 0.25Hz were plotted (thick color lines). Color shaded areas represent standard error of the mean. Coherence between two time series at the phase lag of 0±π/2 indicates positive correlation, while negative correlation is represented by the coherence at the phase lag of π±π/2. The mean time-averaged coherence between RVT and ΔBOLD in all three brain regions consistently increased at the frequency range from 0.016 to 0.031Hz at the phase lag of 0±π/2, while the mean time-averaged coherence at the phase lag of π±π/2 increased from 0.008 to

0.016Hz except in the right IPL. Such changes in the time-averaged coherence between RVT and ΔBOLD are different from those between RGE metrics and ΔBOLD.

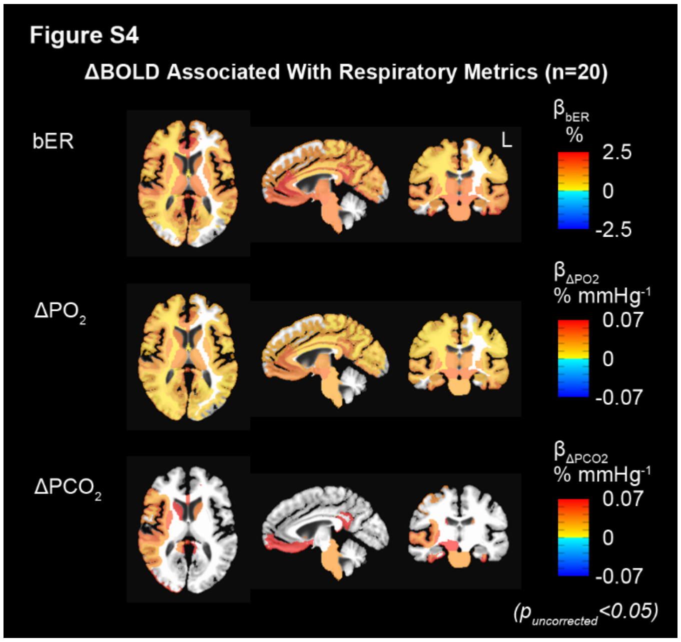


Figure S4. Regional association between ΔBOLD and RGE metrics for all subjects included in the MRI sessions (n=20) before correcting for multiple comparisons. Group maps showing significant changes of regional $\beta_{bER}$ and $\beta_{\Delta PO2}$ before correcting for multiple comparisons were similar to those after correcting for multiple comparisons in Figure 4. Brain regions showing changes of $\beta_{\Delta PCO2}$ before correction could not survive after correction in Figure 4.

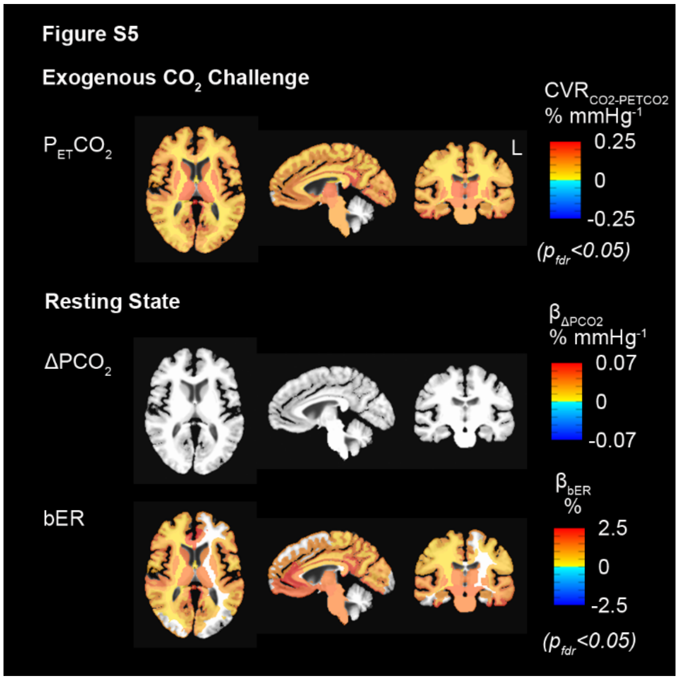


Figure S5. Group CVR showing significant CVR changes under exogenous $CO_2$ challenge (n=10) and during spontaneous breathing at rest (n=20). False discovery rate was used to correct for multiple comparisons and significant association/differences were considered at false discovery rate adjusted $p_{fdr}<0.05$. The CVR map during spontaneous breathing indicated by $\beta_{bER}$ changes resembled the CVR map under exogenous $CO_2$ challenge indicated by $CVR_{CO2\text{-}PETCO2}$.

| | TCD | | | BOLD | | |
|---|---|---|---|---|---|---|
| **Subjects** | **$\Delta PO_2$ vs $\Delta PCO_2$** | **bER vs $\Delta PO_2$** | **bER vs $\Delta PCO_2$** | **$\Delta PO_2$ vs $\Delta PCO_2$** | **bER vs $\Delta PO_2$** | **bER vs $\Delta PCO_2$** |
| s1 | --- | --- | --- | 0.672 (<0.001) | 0.895 (<0.001) | 0.272 (<0.001) |
| s2 | --- | --- | --- | 0.437 (<0.001) | 0.280 (0.002) | -0.740 (<0.001) |
| s3 | --- | --- | --- | 0.191 (0.085) | 0.379 (<0.001) | -0.830 (<0.001) |
| s4 | --- | --- | --- | 0.774 (<0.001) | 0.860 (<0.001) | 0.343 (<0.001) |
| s5 | --- | --- | --- | 0.667 (<0.001) | 0.678 (<0.001) | -0.043 (0.556) |
| s6 | --- | --- | --- | 0.896 (<0.001) | 0.901 (<0.001) | 0.633 (<0.001) |
| s7 | --- | --- | --- | 0.956 (<0.001) | 0.994 (<0.001) | 0.921 (<0.001) |
| s8 | --- | --- | --- | 0.778 (<0.001) | 0.728 (<0.001) | 0.137 (0.100) |
| s9 | --- | --- | --- | 0.419 (<0.001) | 0.764 (<0.001) | -0.261 (0.003) |
| s10 | 0.844 (<0.001) | 0.934 (<0.001) | 0.598 (<0.001) | 0.814 (<0.001) | 0.907 (<0.001) | 0.495 (<0.001) |
| s11 | 0.807 (<0.001) | 0.726 (<0.001) | 0.180 (0.026) | 0.804 (<0.001) | 0.869 (<0.001) | 0.419 (<0.001) |
| s12 | 0.938 (<0.001) | 0.969 (<0.001) | 0.833 (<0.001) | 0.852 (<0.001) | 0.844 (<0.001) | 0.441 (<0.001) |
| s13 | 0.731 (<0.001) | 0.929 (<0.001) | 0.427 (<0.001) | 0.846 (<0.001) | 0.985 (<0.001) | 0.740 (<0.001) |
| s14 | 0.825 (<0.001) | 0.846 (<0.001) | 0.399 (<0.001) | 0.799 (<0.001) | 0.894 (<0.001) | 0.446 (<0.001) |
| s15 | 0.878 (<0.001) | 0.526 (<0.001) | 0.057 (0.518) | 0.845 (<0.001) | 0.733 (<0.001) | 0.256 (0.001) |
| s16 | 0.879 (<0.001) | 0.736 (<0.001) | 0.327 (<0.001) | 0.528 (<0.001) | 0.879 (<0.001) | 0.062 (0.416) |
| s17 | 0.809 (<0.001) | 0.937 (<0.001) | 0.552 (<0.001) | 0.830 (<0.001) | 0.939 (<0.001) | 0.588 (<0.001) |
| s18 | 0.766 (<0.001) | 0.971 (<0.001) | 0.593 (<0.001) | 0.798 (<0.001) | 0.972 (<0.001) | 0.644 (<0.001) |
| s19 | 0.891 (<0.001) | 0.960 (<0.001) | 0.729 (<0.001) | 0.460 (<0.001) | 0.840 (<0.001) | -0.094 (0.217) |
| s20 | 0.752 (<0.001) | 0.572 (<0.001) | -0.101 (0.188) | 0.785 (<0.001) | 0.849 (<0.001) | 0.342 (<0.001) |
| s21 | 0.932 (<0.001) | 0.750 (<0.001) | 0.488 (<0.001) | --- | --- | --- |
| s22 | 0.895 (<0.001) | 0.752 (<0.001) | 0.382 (<0.001) | --- | --- | --- |

Table S1. Strength of correlation indicated by Pearson's correlation coefficients among RGE metrics including bER, $\Delta PO_2$ and $\Delta PCO_2$ in all subjects who participated in TCD sessions (n=13), and those who participated in MRI sessions (n=20). The time series of bER had stronger correlation with that of $\Delta PO_2$ than $\Delta PCO_2$, although both $\Delta PO_2$ and $\Delta PCO_2$ contributed to changes of bER. The correlation coefficients from $\Delta PO_2$ vs $\Delta PCO_2$ varied from 0.7 to 0.9 in TCD sessions and from 0.2 to 0.9 in MRI sessions, suggesting that $\Delta PO_2$ than $\Delta PCO_2$ are not necessarily redundant.

| | ΔCBFv in LMCA | | | | ΔCBFv in RMCA | | | |
|---|---|---|---|---|---|---|---|---|
| **Subjects** | **bER** | **$\Delta PO_2$** | **$\Delta PCO_2$** | **$P_{ET}CO_2$** | **bER** | **$\Delta PO_2$** | **$\Delta PCO_2$** | **$P_{ET}CO_2$** |
| s10 | 0.400 (<0.001) | 0.423 (<0.001) | 0.367 (<0.001) | 0.368 (<0.001) | 0.678 (<0.001) | 0.628 (<0.001) | 0.394 (<0.001) | 0.399 (<0.001) |
| s11 | 0.525 (<0.001) | 0.406 (<0.001) | 0.126 (0.121) | 0.159 (0.051) | 0.207 (0.011) | 0.243 (0.003) | 0.167 (0.040) | 0.179 (0.027) |
| s12 | 0.858 (<0.001) | 0.849 (<0.001) | 0.752 (<0.001) | 0.776 (<0.001) | 0.877 (<0.001) | 0.854 (<0.001) | 0.736 (<0.001) | 0.763 (<0.001) |
| s13 | 0.505 (<0.001) | 0.400 (<0.001) | 0.052 (0.521) | 0.082 (0.313) | 0.379 (<0.001) | 0.380 (<0.001) | 0.231 (0.004) | 0.234 (0.004) |
| s14 | 0.459 (<0.001) | 0.464 (<0.001) | 0.313 (<0.001) | 0.370 (<0.001) | 0.362 (<0.001) | 0.289 (<0.001) | 0.121 (0.104) | 0.130 (0.080) |
| s15 | 0.295 (0.001) | 0.301 (<0.001) | 0.188 (0.032) | 0.193 (0.027) | 0.461 (<0.001) | 0.325 (<0.001) | 0.113 (0.199) | 0.124 (0.159) |
| s16 | 0.571 (<0.001) | 0.314 (<0.001) | 0.039 (0.656) | 0.032 (0.721) | 0.471 (<0.001) | 0.254 (0.003) | 0.019 (0.826) | 0.017 (0.850) |
| s17 | --- | --- | --- | --- | 0.293 (<0.001) | 0.356 (<0.001) | 0.359 (<0.001) | 0.359 (<0.001) |
| s18 | 0.780 (<0.001) | 0.762 (<0.001) | 0.475 (<0.001) | 0.483 (<0.001) | 0.758 (<0.001) | 0.755 (<0.001) | 0.503 (<0.001) | 0.501 (<0.001) |
| s19 | 0.723 (<0.001) | 0.724 (<0.001) | 0.599 (<0.001) | 0.584 (<0.001) | 0.764 (<0.001) | 0.767 (<0.001) | 0.646 (<0.001) | 0.638 (<0.001) |
| s20 | 0.426 (<0.001) | 0.469 (<0.001) | 0.235 (0.002) | 0.167 (0.028) | 0.439 (<0.001) | 0.573 (<0.001) | 0.344 (<0.001) | 0.271 (<0.001) |
| s21 | 0.582 (<0.001) | 0.278 (<0.001) | 0.048 (0.522) | 0.110 (0.136) | --- | --- | --- | --- |
| s22 | 0.509 (<0.001) | 0.286 (<0.001) | 0.065 (0.374) | 0.082 (0.260) | 0.470 (<0.001) | 0.214 (0.003) | -0.004 (0.951) | 0.022 (0.763) |
| Mean Fisher Z | 0.662 (---) | 0.558 (0.034) | 0.305 (<0.001) | 0.320 (<0.001) | 0.622 (---) | 0.566 (0.175) | 0.339 (<0.001) | 0.342 (<0.001) |

Table S2. Strength of correlation indicated by Pearson's correlation coefficients between ΔCBFv and RGE metrics including bER, $\Delta PO_2$, $\Delta PCO_2$ and $P_{ET}CO_2$. Numbers in brackets next to Pearson's correlation coefficients indicate p values from individual correlation analyses. The bottom row shows the mean values of Fisher Z scores transformed from Pearson's correlation coefficients in groups. Numbers in brackets next to mean Fisher Z scores indicate p values in the paired comparisons. The correlation between ΔCBFv and bER was significantly stronger than the correlation between ΔCBFv and $\Delta PCO_2$/$P_{ET}CO_2$ in the paired comparisons ($p<0.001$). bER and $\Delta PO_2$ are the parameters that consistently showed significantly high correlation with the ΔCBFv measured in LMCA and RMCA.
